# CRISPR-Cas supervises diverse anti-phage defense systems

**DOI:** 10.1101/2024.12.27.630466

**Authors:** Xian Shu, Rui Wang, Feiyue Cheng, Zhihua Li, Aici Wu, Qiong Xue, Chao Liu, Xifeng Cao, Yue Feng, Ming Li

## Abstract

A variety of bacterial anti-phage systems have recently been discovered^1–3^, but how these systems synergize to defend against diverse phages remains poorly understood. Here, we report that the adaptive immune system CRISPR-Cas supervises the expression of diverse immune systems by exploiting the regulatory CRISPR RNA-like RNAs (crlRNAs). The crlRNAs target and inhibit the promoters of various immune systems, including the newly characterized Nezha and Gabija, as well as eight previously unrecognized systems that feature distinct defensive domains. Notably, CRISPR regulation balances the expression level of these systems to ensure effective anti-phage activity while avoiding their autoimmunity risks. In return, the supervised immune systems trigger abortive infections when CRISPR-Cas is inhibited by viral anti-CRISPR proteins, thereby offering an anti-anti-CRISPR protection at the population level. Moreover, these systems complement CRISPR immunity with a differing anti-phage profile. These findings highlight the pivotal role of CRISPR-Cas in orchestrating a diverse range of immune systems and showcase the delicate synergy among the multilayered defense strategies in prokaryotes.

## Main

CRISPR (Clustered Regularly Interspaced Short Palindromic Repeat) arrays, in conjunction with Cas (CRISPR-associated) proteins offer prokaryotes an adaptive immune mechanism against invading mobile genetic elements (MGEs) like plasmids and bacteriophages^4,5^. The CRISPR RNAs (crRNAs) contain information from MGEs, guiding Cas proteins to specifically target and eliminate the foreign DNA/RNA. Similar to other immune systems, CRISPR-Cas often co-exist with diverse defensive genes or systems in bacterial genome, forming defense islands that are believed to spread as a whole throughout the bacterial population^6,7^. This genetic linkage has led to the discovery of numerous new immune systems in recent years^1–3^. However, how CRISPR-Cas interplays and synergizes with these systems to defend host cells against various phages remains poorly understood. Here, we demonstrate that CRISPR-Cas supervises the expression of a range of coexisting immune systems, many of which were previously unknown, working together to form a reciprocal and synergistic ‘anti-phage’ combination.

### Transcriptional regulation of Nezha and Gabija by I-C CRISPR-Cas

Recent studies have revealed that CRISPR-Cas systems also play a role in gene regulation, in addition to their canonical role in adaptive immunity. Some solitary CRISPR repeat units (SRUs) produce crRNA-like RNAs (crlRNAs) that target the promoter regions of endogenous genes (such as toxin genes) to modulate their expression in a Cas-dependent manner, notably without incurring DNA cleavage due to the limited base pairings^8–10^. In our search for crlRNAs associating with type I-C systems, we observed that a Nezha system is encoded within the *cas* operon of 12 bacterial strains, predominantly from *Neisseria* species (Supplementary Data 1). The Nezha system consists of a sirtuin (Sir2) NADase and a HerA helicase, which together form a supramolecular octadecamer with multiple enzymatic activities involved in phage defense^11,12^. Among the 12 Nezha-associated *cas* operons, an SRU is consistently located between the *herA* and *cas5c* genes (Fig. 1a and Supplementary Data 1). Our small RNA sequencing (sRNA-seq) data confirmed that the SRU from *Neisseria subflava* CCUG 29761 produced crlRNAs, typically 48-50 nt in length, in *E. coli* cells encoding their corresponding Cascade proteins (Cas5c, Cas8c, Cas7c, and Cas11) (Fig. 1a). Notably, similar to a typical CRISPR repeat RNA processed by the Cas5c endonuclease^13^, the 5′ repeat-like (ΨR) sequence of crlRNA precursor was cleaved 3′ directly of a hairpin structure and 5′ of a conserved 11 nt segment (retained on mature crlRNAs as a 5′-handle). In contrast, the 3′ end of the crlRNAs terminates variably within a loop of a 3′ hairpin structure (Fig. 1a), suggesting a different processing mechanism (see below).

**Figure 1:**
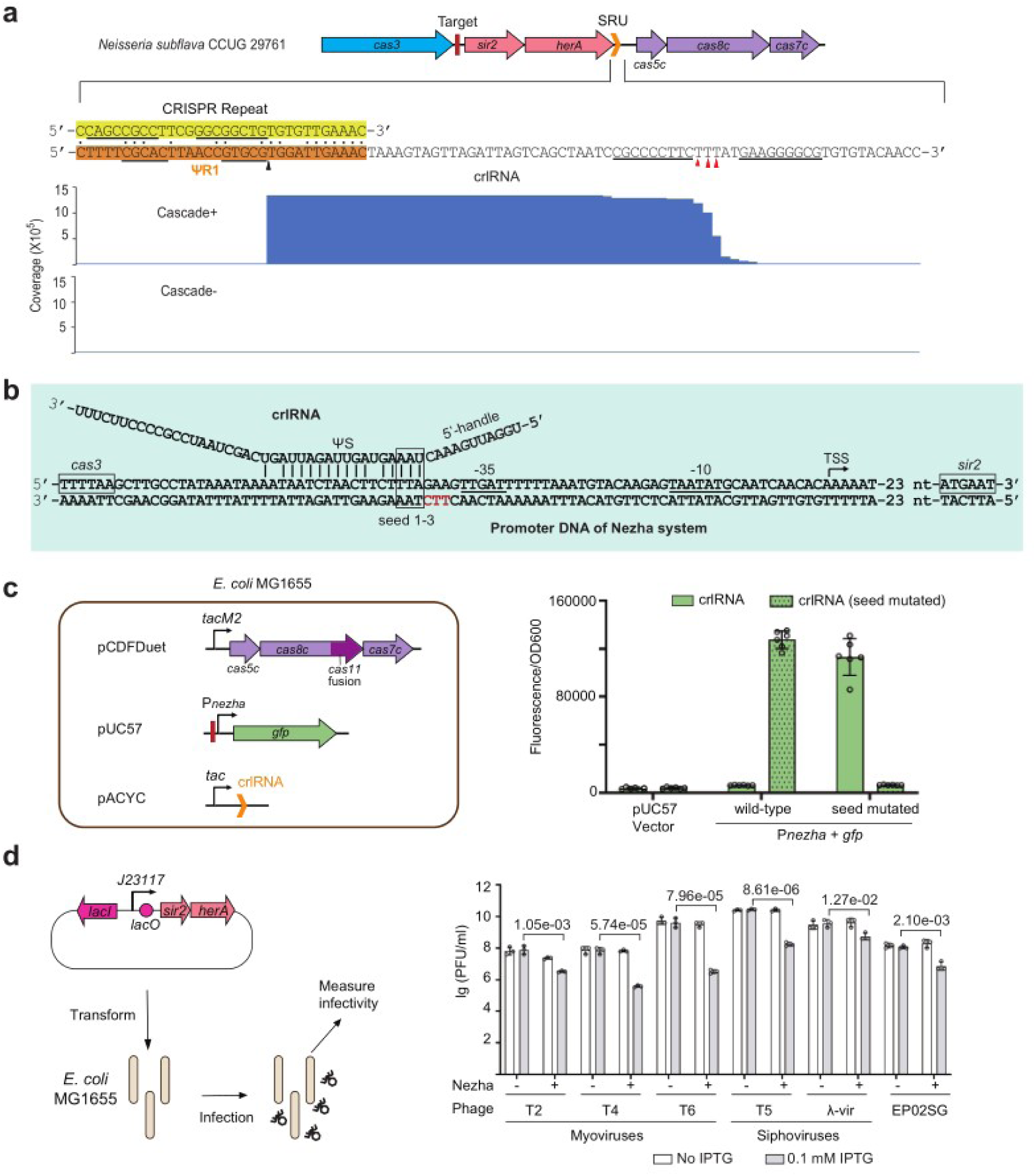
A Nezha system regulated by type I-C CRISPR-Cas. **a**, Genetic organization of a CRISPR-regulated Nezha system and determination of the regulatory crlRNA. The 5′ (black triangle) and 3′ (red triangles) termini of crlRNA were determined by sRNA-seq in *E. coli* cells expressing (Cascade+) or lacking (Cascade-) the Cascade proteins. Dots indicate identical nucleotides shared between the CRISPR repeat and the repeat-like (ΨR) sequence of crlRNA. Palindromic nucleotides forming the stem of an RNA hairpin structure are underlined. **b**, The complementarity between crlRNA and its target site within the promoter of Nezha (P*_nezha_*). TSS, transcription start site (determined by primer extension; see Extended Data Fig. 1). PAM nucleotides are highlighted in red. Framed complementing nucleotides were subjected to mutation analysis in panel **c**. **c**, Fluorescence from a P*_nezha_*-controlled green fluorescence protein (*gfp*) gene in the presence of the corresponding crlRNA and Cas proteins. The red bar indicates the target site of crlRNA. *tacM2*, a mutated version of *tac* promoter. The *cas11* gene is fused within *cas8c*. Data are presented as the mean ± s.d. from six biologically independent replicates. **d**, Plaque forming units of *E. coli* phages infecting MG1655 cells expressing the Nezha system (controlled by an inducible *J23117* promoter). Data are presented as the mean ± s.d. from three biologically independent replicates. EP02SG is a non-model phage isolated from sewage water, with its genome information available in NCBI (1181).

Importantly, the spacer-like (ΨS) sequence of crlRNA shows a 16-bp complementarity (with a mismatch at position 6) to the promoter of Nezha (P*_nezha_*) along with a 5′-TTC-3′ protospacer adjacent motif (PAM) (Fig. 1b). By characterizing P*_nezha_* using primer extension (Extended Data Fig.1), we noticed that its -35 element locates adjacent to the target site of crlRNA (Fig. 1b). In *E. coli* cells expressing the corresponding Cascade proteins, the crlRNA markedly inhibited a P*_nezha_*-controlled green fluorescence gene (*gfp*), with a suppressive factor of 21.3 (compared to a mutated crlRNA lacking complementarity to P*_nezha_*) (Fig. 1c). Notably, when a complementarily mutated P*_nezha_* was employed to drive *gfp* transcription, the mutated crlRNA showed inhibition instead of the wild type (Fig. 1c). We conclude that crlRNAs direct Cas proteins to inhibit the promoter of the associated Nezha system. Using a series of *E. coli* phages, we validated that this unique Nezha system effectively defends *E. coli* cells against T2, T4, T5, T6, and virulent λ phage infections (Fig. 1d).

The anti-phage defense system Gabija comprises of an OLD family nuclease, GajA, and a UvrD-like helicase, GajB^14–16^. Similarly, we discovered Gabija-associated type I-C CRISPR-Cas loci in four *Neisseria* strains (Extended Data Fig. 2a and Supplementary Data 1). We have also detected highly conserved SRUs between their *gajB* and *cas5c*, with their target sites consistently positioned upstream of *gajA* (Extended Data Fig. 2a). Using the *gfp* reporter gene, we demonstrated that the Gabija promoter (P*_gaj_*) is inhibited by the conserved SRU by a factor of 22.4 in *E. coli* cells expressing type I-C Cascade proteins (Extended Data Fig. 2c). Notably, unlike the SRUs governing Nezha, the Gabija-regulating SRUs each contain two sequences closely resembling CRISPR repeat (designated as ΨR1 and ΨR2), and in line with a typical I-C crRNA, their primary crlRNA products (57 nt) featured an 11-nt 5′ handle and a 21-nt 3′ handle, albeit alternative/subsequent processing of their 3′ end was less frequently observed (Extended Data Fig. 2b).

### Exploring new immune systems based on CRISPR regulation

Motivated by the above findings, we sought out to explore other immune systems regulated by type I-C CRISPR-Cas. By scanning the genetic sequence between *cas3* and *cas5c*, we initially identified a group of I-C *cas* operons that are intervened by various non-Cas genes. Subsequently, we searched for SRUs and their target sites within the intergenic regions. As a result, we obtained eight previously unrecognized candidate immune systems likely controlled by CRISPR-Cas (Fig. 2a and Supplementary Data 1), and collectively referred to them as CRISIS (<u>CRI</u>SPR-<u>s</u>upervised immune <u>s</u>ystem) for convenience. Among these systems, one encodes a small standalone Sir2-like protein, so we named this system sole-Sir2. To differentiate the other seven systems, we named them after Chinese deities (Fig. 2a). The genes of each CRISIS are directly followed by an SRU (preceding *cas5c*), which partially complements to a target sequence right preceding CRISIS genes (with a 5′-TTC-3′ or 5′-TTA-3′ PAM) (Fig. 2a). We employed sRNA-seq to analyze the RNA products from five SRUs, all of which were predominantly 49-50 nt in length, featuring a conserved 11-nt 5′ handle and a flexible 3′ terminus, akin to the crlRNA regulating Nezha (Extended Data Fig. 3). Notably, these 3′ termini were found to be positioned conservatively within the loop of a 3′ hairpin structure. In contrast, our *in vitro* experiments observed that Cas5c generated a canonical cleavage directly 3′ of these hairpin structures (Extended Data Fig. 4), indicating that noncanonical or subsequent 3′ processing should have occurred *in vivo*.

**Figure 2:**
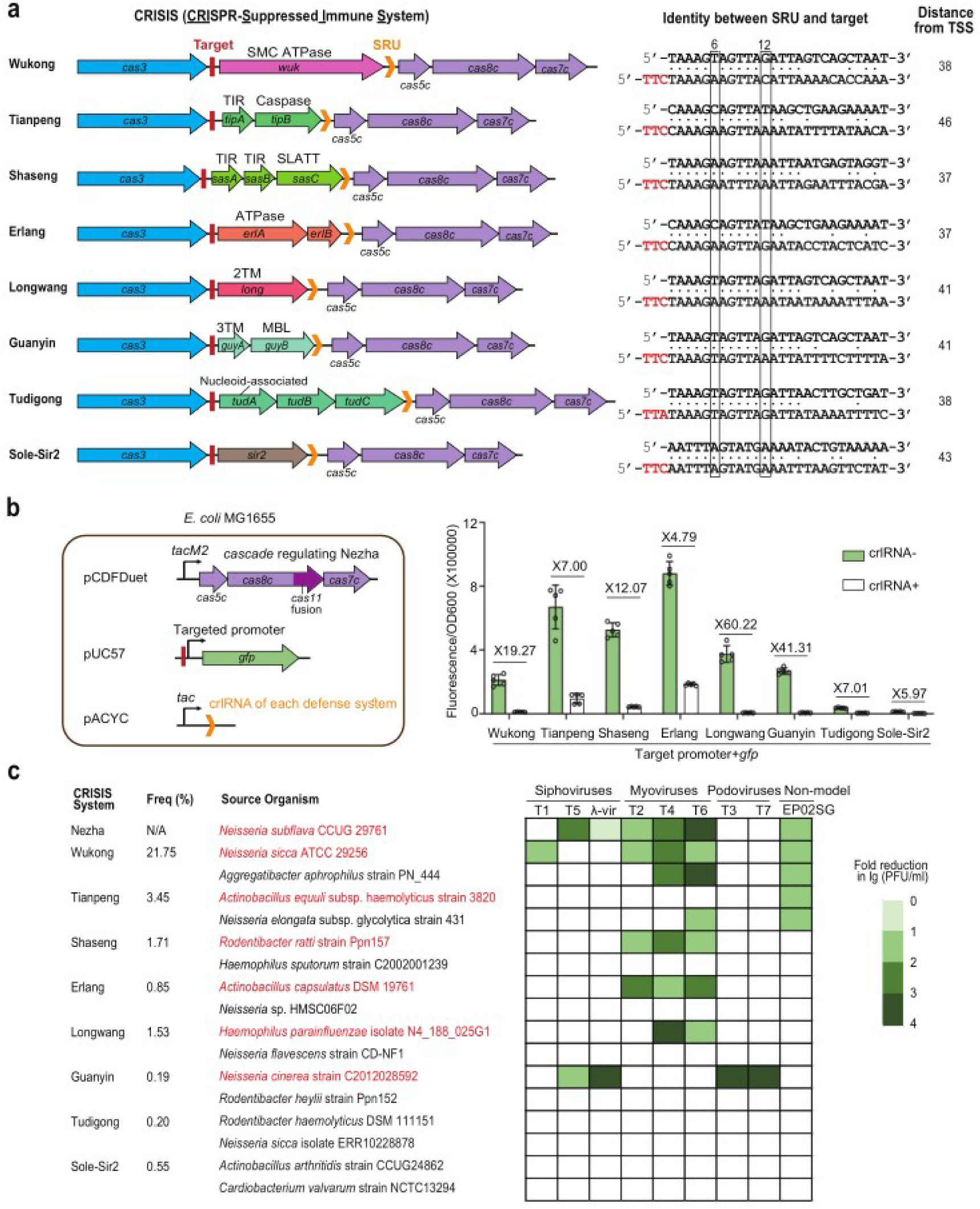
More CRISPR-supervised immune systems (CRISIS). **a**, Genetic architecture of 8 CRISIS systems (each named after a Chinese deity). For each CRISIS, the nucleotide identity between a representative SRU and its target is given. The indicated 6^th^ and 12^th^ nucleotide positions do not participate in guide-target base pairing (Zhao et al., 2014). PAM nucleotides are highlighted in red with their distance (nt) from the TSS of CRISIS genes given. To avoid distractions, *cas* genes involved in adaptation are not depicted. **b**, Fluorescence from a *gfp* gene controlled by the promoter of different CRISIS systems in the presence or absence of the corresponding crlRNA. Note that *E. coli* cells expressing the *N. subflava* Cas proteins regulating Nezha were used for this assay. Data are presented as the mean ± s.d. from six biologically independent replicates. **c**, Immunity of each CRISIS against different *E. coli* phages infecting MG1655 cells. See detailed information in Extended Data Fig. 5. The frequency of bacterial genomes predicted to contain each CRISIS system is given. Organism names in red indicate the source of CRISIS systems further characterized in Figure 3-5.

Interestingly, mismatches between the SRUs and their target tended to occur at the PAM-distal region or at the 6^th^ and 12^th^ positions (Fig. 2a), which do not participate in base pairing during target DNA recognition^17^. Through primer extension, we identified that the transcription start site (TSS) of each CRISIS lies 37-46 bp downstream of the target site of corresponding SRUs (Fig. 2a), suggesting that Cascade-crlRNA binding to these target sites may interfere with the function of their promoter elements. In *E. coli* cells expressing the Cascade proteins known to regulate Nezha, we measured the strength of the promoters of these CRISIS systems in the presence or absence of the corresponding crlRNA (Fig. 2b). As expected, the promoter activity of each CRISIS was inhibited by crlRNA, with inhibition factors ranging from 4.79 to 60.22.

Next, we assessed the immune effects of these CRISIS systems in *E. coli* cells. For each CRISIS, we synthesized its encoding sequences from two or more bacterial organisms (Fig. 2c) and introduced them into MG1655 cells under an inducible promoter. For most systems (except Tudigong and sole-Sir2), we observed immunity against at least one specific *E. coli* phage (Fig. 2c and Extended Data Fig. 5). Notably, the same CRISIS system derived from different organisms exhibited varying anti-phage profiles, indicating their specificity in sensing and/or responding to phage infections. This specificity may have hindered us from elucidating the immune effects of Tudigong and sole-Sir2 with our current phage collection. In accord, we were able to identify a close homolog of sole-Sir2, lacking association with CRISPR-Cas, that effectively protected *E. coli* cells against T2 and T4 phages. (Extended Data Fig. 6a).

### CRISIS systems exploit diverse enzymatic activities

We then looked into the functional domains of each CRISIS system. The sole-Sir2 system encodes a single ∼440 amino acid (aa) protein (Fig. 2a), which contains the Sir2-like domain capable of degrading nicotinamide adenine dinucleotide (NAD+)^12^ (Extended Data Fig. 6b). Consistently, sole-Sir2 was active in converting NAD+ to ADP-ribose (ADPR) (Extended Data Fig. 6c). Structural prediction and alignment revealed that its C-terminal Sir2 domain shares significant similarity with the N-terminal of a recently characterized large Defense-associated sirtuin 2 (DSR2) protein (∼943 aa)^12,18^, while its N-terminal lacks structural resemblance to DSR2 (Extended Data Fig. 6d). We speculate that sole-Sir2 may sense phage infections through this differing N-terminal substructure.

Wukong encodes a single large DNase (∼1026 aa) containing an N-terminal PHP exonuclease domain and a C-terminal ATPase domain, and actively degraded various DNA substrates *in vitro* (Fig. 3a). As expected, mutating the active site of either domain resulted in the inactivation of this immune system (Fig. 3a). In comparison, Erlang encodes a smaller ATPase (ErlA; ∼458 aa) along with a hypothetical protein (ErlB), and mutating the predicted active site of ErlA disabled this immune system (Fig. 3b). Interestingly, co-expression of ErlA and ErlB led to the formation of a ∼300 kDa complex, likely comprising 4 ErlA and 4 ErlB subunits (Extended Data Fig. 7a-c), which degraded the genomic DNA of T4 and T6 phages *in vitro* (Fig. 3b).

**Figure 3:**
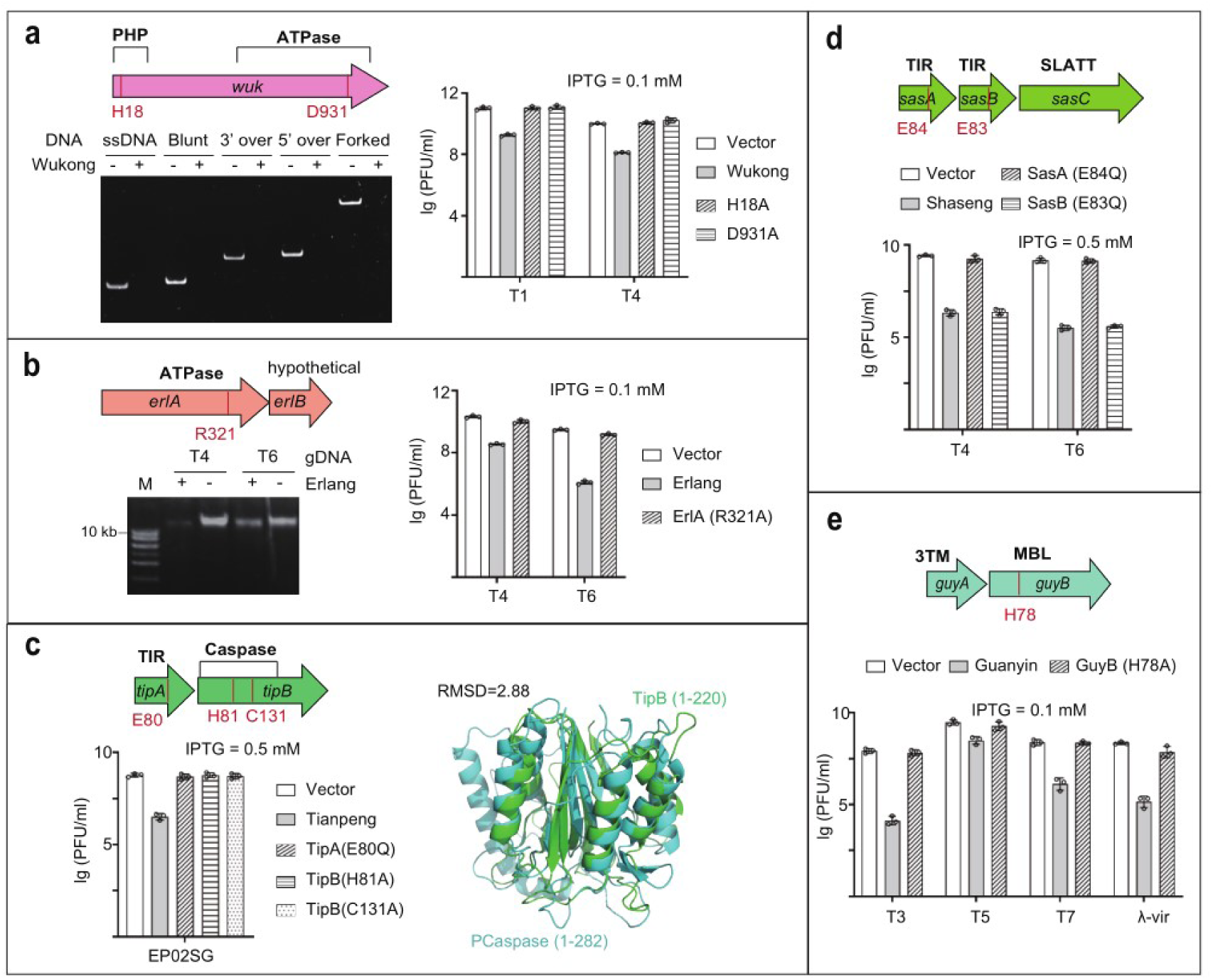
Diverse functional domains of the CRISIS systems. **a**-**b**, Characterization of the DNase activity of Wukong (**a**) and Erlang (**b**) systems. For Wukong, various DNA substrates were tested, including single-stranded DNA (ssDNA), double-stranded DNA with blunted ends (blunt), 3′ or 5′ overhangs, and forked DNA. For the Erlang complex (see Extended Data Fig. 7a-c), the genomic DNA (gDNA) of T4 or T6 was used as substrate. M, 1-kb DNA ladder. **c-d**, Mutation analysis of the TIR and effector proteins of Tianpeng (**c**) and Shaseng (**d**). For Tianpeng, structural alignment of the N-terminal substructures from TipB and the PCaspase protein associated with type III-B CRISPR-Cas (Jurre Steens, et al., 2024) is provided. **e**, Mutation analysis of the MBL metallo-hydrolase from Guanyin. This system also encodes a 3-transmembrane (3TM) protein (GuyA). Data are presented as the mean ± s.d. from three biologically independent replicates.

Tianpeng and Shaseng encode a Caspase protease and a SLATT transmembrane (TM) protein, respectively, associating with one or two proteins containing the Toll/interleukin-1 receptor (TIR) domain (Fig. 3 c and d). The TIR domain is a conserved component among eukaryotic and prokaryotic immune systems^19,20^, and in the bacterial Thoeris system, it plays a crucial role in sensing phage infection and producing signaling molecules to activate either a Sir2 NADase or a TM protein^21^.

Notably, these Thoeris effectors bind to the signaling molecule through a C-terminal SLOG or MACRO domain, which are absent in Caspase and SLATT proteins. This suggests that Caspase and SLATT likely sense the TIR-produced signaling molecules through distinct mechanisms. Consistently, inactivating the TIR protein of each system abolished their immune responses (Fig. 3 c and d). Interestingly, though Shaseng encodes two TIR proteins, only dead mutation of SasA abolished phage immunity (Fig. 3d). We speculate SasB may interact with SasA and assist it in sensing phage components, as suggested in a recent preprint about a staphylococcal two-TIR Thoeris defense system^22^. Additionally, we observed that the N-terminal of the Caspase protein TipB shares a high structural resemblance (RMSD=2.88) with the N terminal of the prokaryotic Caspase (PCaspase) (Fig. 3c). PCaspase is cleaved and activated by the SAVED-CHAT protease, which senses the cyclic tri-adenosine monophosphate produced from type III-B CRISPR-Cas systems^23^. Based on the structural alignment, we predicted two potential active sites (His81 and Cys131) of TipB, and separate mutations of these sites inactivated the Tianpeng system (Fig. 3c).

Longwang encodes a single 2TM protein, suggesting it anchors on the cell envelope to impart immune responses. Interestingly, its predicted structure features a long α-helix (with a high confidence) that separates the 2TM domain from a four-helix substructure (Extended Data Fig. 7d). Guanyin similarly encodes a 3TM protein (GuyA), as well as an MBL metallo-hydrolase (GuyB) that closely resembles the binuclear Zn^2+^ catalytic module of the teichoic acid phosphorylcholine esterase (Pce)^24^ (RMSD=3.62; Extended Data Fig. 7e). Consistently, mutating the predicted catalytic residue (His78) of GuyB resulted in the inactivation of this immune system (Fig. 3e). Notably, GuyB lacks the choline-binding module of Pce (Extended Data Fig. 7e), suggesting that it may break phosphate ester bonds of other essential molecules in the cell. At last, Tudigong encodes two hypothetical proteins along with a nucleotide-associating protein that putatively interacts directly with phage DNA (Fig. 2a).

Overall, CRISIS systems employ various functional domains to achieve immunity against phages. Preliminary bioinformatic analyses showed that these immune systems (which not necessarily co-occur with a CRISPR-Cas locus) are present in bacterial genomes at an estimated frequency ranging from 0.19-21.75% (Fig. 2a). We also conducted phage infection assays at different multiplicity of infections (MOIs), and found that most of these systems provided effective phage resistance only under low MOI conditions (Extended Data Fig. 8), which suggests the induction of abortive infection (Abi) responses during their immune processes. Further research is required to fully understand their detailed mechanisms in detecting phage infections and initiating immune responses.

### CRISPR supervision ensures CRISIS immunity and prevents its autoimmunity

We then asked whether CRISIS systems could provide effective immunity under the transcriptional suppression by CRISPR-Cas. We selected the Nezha, Wukong, and Shaseng systems, which had previously been shown to restrict T4 and T6 infections (Fig. 2c). For each CRISIS system, we synthesized, codon-optimized, and cloned the original sequences encoding Cas3, CRISIS, and Cascade proteins into a vector (designated as pCas-CRISIS) and introduced them into *E. coli* cells to evaluate their immunity against T4 and T6 (Fig. 4a). Note that CRISPR immunity was excluded in these assays as no phage-targeting crRNAs were provided. Remarkably, pCas-Nezha, pCas-Wukong, and pCas-Shaseng all demonstrated active protection (Fig. 4a), which seemed to be more effective compared to their performance in Fig. 2c and Extended Data Fig. 5. For example, pCas-Nezha and pCas-Wukong provided nearly complete resistance to T4 and T6, making it difficult to calculate PFU (Fig. 4a), whereas only a 2-3 log reduction in PFU was observed in Extended Data Fig. 5. Notably, when start codon mutations were introduced into the CRISIS genes, phage immunity was no longer observed (Fig. 4a). Therefore, CRISPR regulation allows for an expression level of CRISIS systems that is sufficient to produce effective anti-phage immunity.

**Figure 4:**
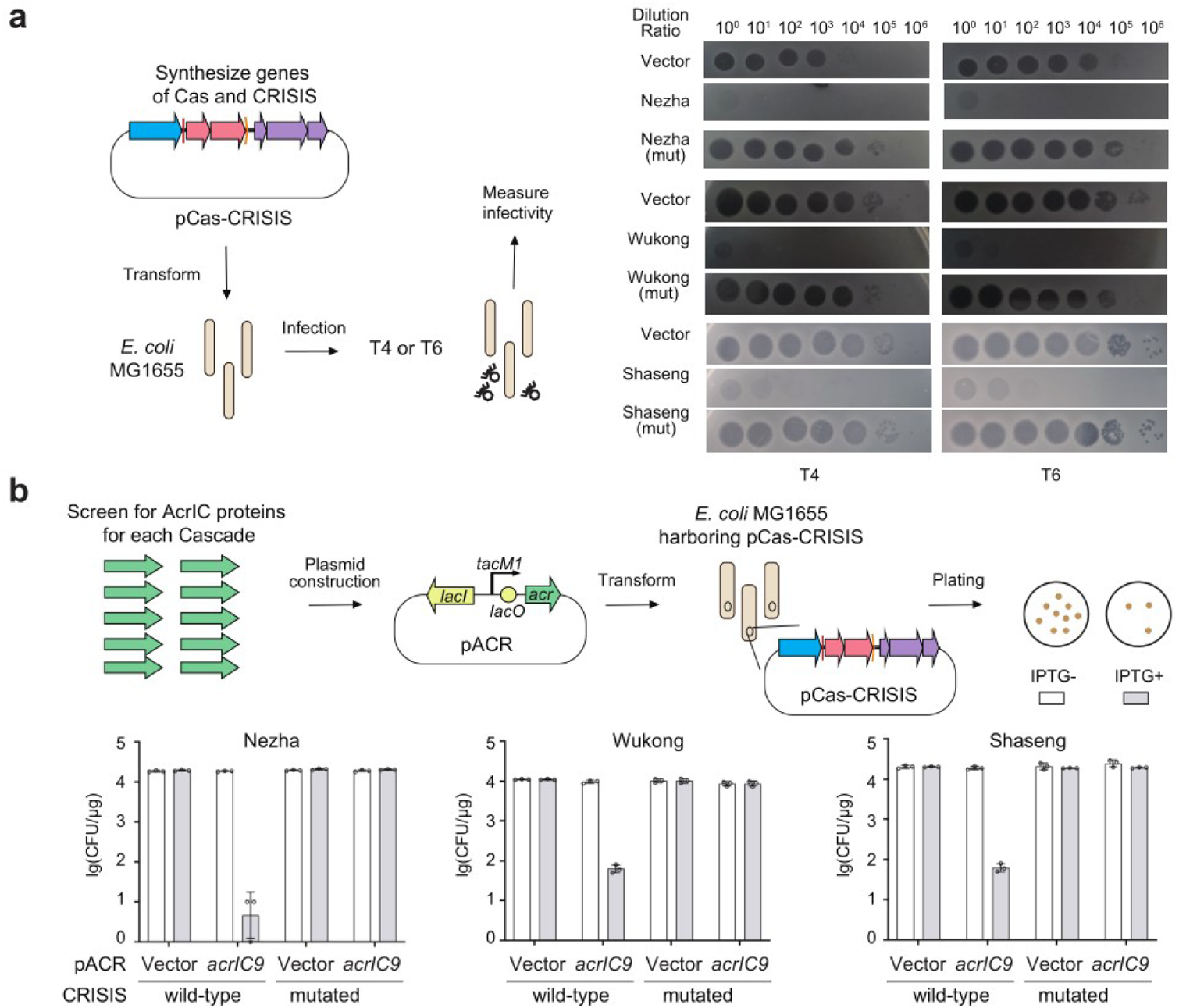
CRISIS immunity and autoimmunity under CRISPR supervision. **a**, The immunity of Nezha, Wukong, and Shaseng against T4 or T6 under the natural transcriptional suppression by the co-occurring Cas proteins. pCas-CRISIS carries the original sequence organization of *cas3*, CRISIS genes, and the Cascade genes. For each CRISIS, a mutant (the start codon of the first gene was mutated) was included as a control. **b,** Transformation efficiency of cells containing pCas-CRISIS with a plasmid expressing AcrIC9 that specifically inhibits the Cascade proteins regulating Nezha, Wukong, and Shaseng (see Extended Data Fig. 10). Acr expression was controlled by a mutated IPTG-inducible *tac* promoter (*tacM1*). Data are presented as the mean ± s.d. from three biologically independent replicates.

We then examined the physiological significance of CRISPR regulation on CRISIS systems. When strong inducible promoters were used to control CRISIS expression, we observed detrimental effects on the growth of *E. coli* cells (Extended Data Fig. 9). We propose that, without CRISPR regulation, the majority, if not all, CRISIS systems will be cytotoxic under the control of their native promoters. Consistently, attempts to clone most CRISIS systems under their native promoters in the absence of their associated Cas proteins were unsuccessful. To address this issue, we undertook a search for anti-CRISPR (Acr) proteins that could specifically inhibit the Cascade proteins regulating a CRISIS system, using our recently developed strategy^25^. We constructed a *gfp* gene controlled by the promoter of Nezha, Wukong, or Shaseng, which was expected to be repressed in the presence of the corresponding Cascade proteins and crlRNAs (Extended Data Fig. 10). Subsequently, we synthesized a series of Acr proteins and tested their ability to induce fluorescence production by inhibiting Cascade proteins. As a result, we identified that AcrIC9 could effectively activate all the three promoters, indicating its capability of inhibiting their respective Cascade proteins (Extended Data Fig. 10). Then we introduced an AcrIC9-enconding plasmid into *E. coli* cells containing pCas-Nezha, pCas-Wukong, or pCas-Shaseng (Fig. 4b). Remarkably, AcrIC9 expression markedly reduced the transformation efficiency of all these three cell types by approximately 2-4 logs. In contrast, this reduction was no longer observed when CRISIS genes were mutated (Fig. 4b). Therefore, CRISIS systems have the potential to induce cell death or autoimmunity in the loss of CRISPR supervision.

### CRISIS provides anti-anti-CRISPR protection and improves anti-phage profile

We then asked how CRISPR-Cas systems benefit from supervising diverse CRISIS systems. Our recent study unraveled that CRISPR-repressed toxin-antitoxins (CreTA) can be activated to kill the host cell when Cas proteins are inhibited by a variety of Acr proteins, and this activation leads to the elimination of Acr-encoding MGEs from the bacterial population, demonstrating a broad-spectrum anti-anti-CRISPR strategy at the population level^25^. We propose that CRISIS systems also provide anti-anti-CRISPR protection. To test this hypothesis, we constructed two reporter plasmids, one with *gfp* (pGFP) and the other with *mcherry* (pMC), and introduced them in a 1:1 ratio to *E. coli* cells containing pCas-Wukong (Fig. 5a). By measuring the green and red fluorescence levels, we monitored their persistence in the bacterial population. Notably, when pMC expressed AcrIC9 that inactivates the Cas proteins controlling Wukong expression, this plasmid was rapidly eliminated from the bacterial population (Fig. 5a). Likewise, AcrIC9-expressing plasmids were effectively expelled from the population of *E. coli* cells containing pCas-Nezha or pCas-Shaseng (Extended Data Fig. 11). Importantly, this effect was abolished when CRISIS genes were mutated. Therefore, CRISIS systems provide anti-anti-CRISPR protection for CRISPR immunity by eliciting Abi-based herd immunity against Acr-encoding MGEs.

**Figure 5:**
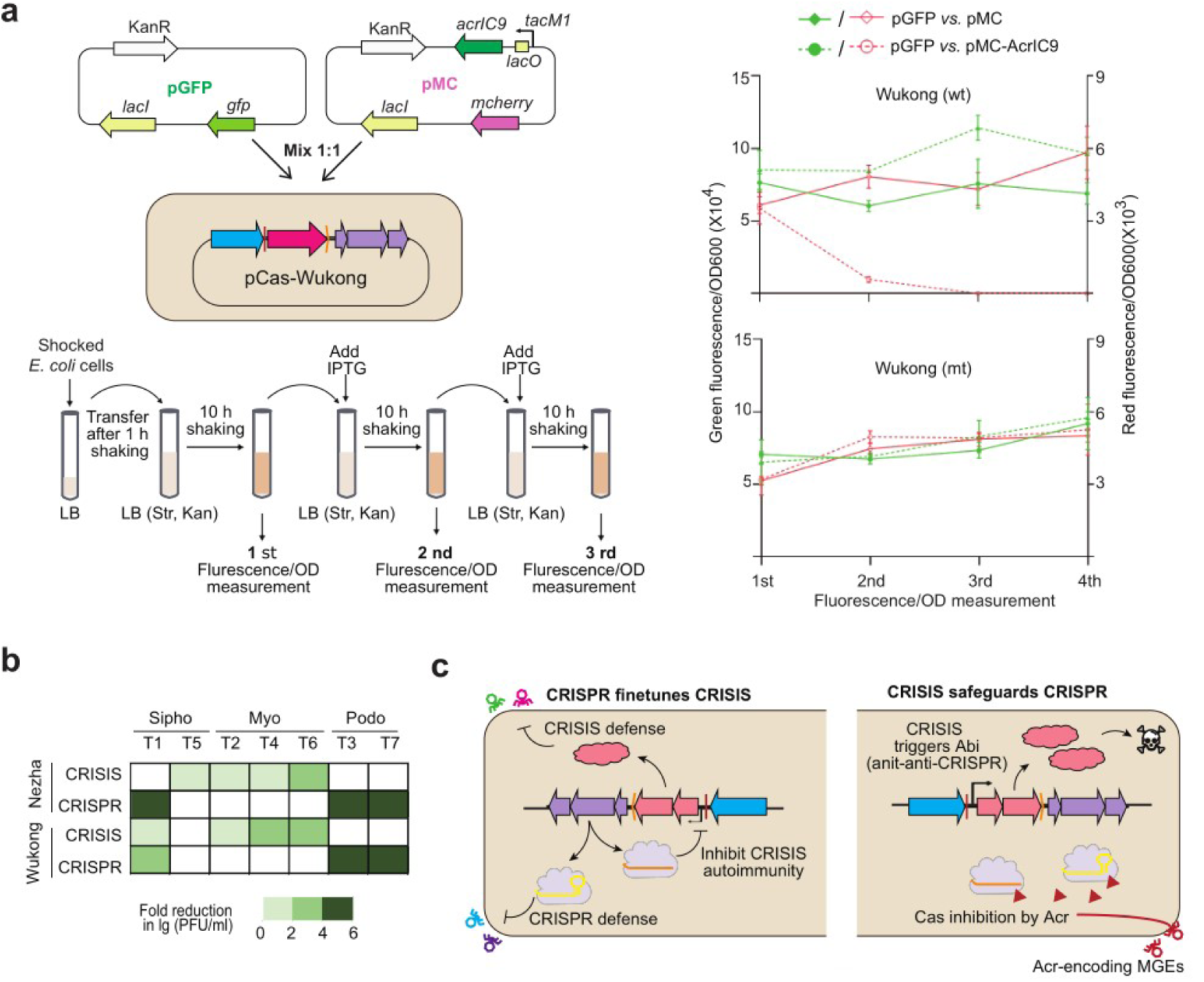
CRISIS safeguards CRISPR-Cas and expands the anti-phage profile. **a**, Persistence of an AcrIC9-encoding plasmid (pMC-AcrIC9; also carrying the *mcherry* reporter gene) in the population of *E. coli* cells containing pCas-Wukong (or a derivate carrying a mutated Wukong system). A competing plasmid (with the same backbone) carrying *gfp* (pGFP) was mixed with pMC-AcrIC9 or pMC (lacking the *acrIC9* gene) at a 1:1 ratio before co-transformation into *E. coli* cells. Data are presented as the mean ± s.d. from three biologically independent replicates. **b**, The differing anti-phage capability between CRISPR-Cas and two CRISIS systems. See detailed data in Extended Data Fig. 12. **c**, The proposed interplay model between CRISPR-Cas and CRISIS systems.

Different defense systems typically have varying anti-phage profiles^26^. Using phage-targeting crRNAs, we demonstrated that Cas proteins regulating Nezha (which can resist T5, T2, T4, and T6; see Fig. 2c) can effectively confer *E. coli* cells with specific CRISPR immunity against T1, T3, and T7, but notably, not against T5, T2, T4, or T6 infections (Fig. 5c and Extended Data Fig. 12). Similarly, Cas proteins monitoring Wukong (resistant to T1, T2, T4, and T6) expression provided CRISPR immunity against T1, T3, and T7. Therefore, the association of CRISPR-Cas loci with diverse CRISIS systems appears to synergistically enhance their anti-phage activity. Intriguingly, type I-C CRISPR-Cas can also regulate the expression of a comprehensive defensive gene cluster (named SMRD), which preferentially encodes a SNF2-related protein, a DNA methyltransferase, a RuhM family nuclease, and a DEAD helicase, along with a known defense system (such as Tudigong, Kiwa, or Erlang) (Extended Data Fig. 13). Thus, it seems that CRISPR-Cas can orchestrate multiple different defense systems to broaden its anti-phage capabilities.

## DISCUSSION

A diverse list of bacterial immune systems has been uncovered in the past six years^1–3^. These anti-phage systems, particularly those that induce Abi responses by targeting essential host components, can become detrimental when they are excessively expressed. It is crucial to carefully regulate the expression of these immune systems to avoid autoimmune effects on the host cell. Otherwise, such effects may lead to the elimination of these immune systems from the rapidly evolving bacterial population. Our research showcases such a regulation mechanism by unravelling several CRISPR-supervised immune systems (CRISIS). CRISPR-based regulation helps in maintaining a balanced expression level of these systems, that ensures effective phage immunity while avoiding potential autoimmune effects (Fig. 4). Remarkably, if the Cas proteins responsible for supervising CRISIS expression are inhibited by Acr proteins, the CRISIS systems become overexpressed to induce an Abi-like response, which can efficiently eliminate Acr-encoding MGEs from the bacterial population, thus providing population-level anti-anti-CRISPR protection for the co-occurring CRISPR-Cas system (Fig. 5a). We propose an interplay model (Fig. 5d) in which CRISPR-Cas systems mitigate the evolutionary disadvantage of various CRISIS systems by lowering their autoimmune risks; in return, the CRISIS systems elevate the evolutionary advantage of CRISPR-Cas by acting as toxic components that need to be inhibited by CRISPR-Cas. This reciprocal relationship fosters a genetic symbiosis of CRISPR-Cas and diverse immune systems. In addition, as CRISPR-Cas and CRISIS systems very likely possess different and complementary anti-phage profiles (Fig. 5c), their combination can result in synergistic and enhanced immunity against phages. A recent study revealed that certain larger type II-C Cas9 proteins are genetically linked with CRISPR-Cas-system-promoting (pro-CRISPR) proteins that promote CRISPR immunity through direct protein interactions^27^. It is conceivable that proteins encoded by some, if not all CRISIS systems, may also directly contribute to CRISPR immunity.

Remarkably, type I-C CRISPR-Cas systems can also regulate the expression of large defensive gene clusters (Extended Data Fig. 13). In addition, we also found Abi-like genes that are possibly regulated by I-B CRISPR-Cas systems, as well as some Fic immune protein encoding genes that are likely controlled by I-F CRISPR-Cas systems (Extended Data Fig. 14 and Supplementary Data 1). Therefore, a common paradigm likely exists among various CRISPR-Cas systems, especially those of type I, in their involvement in orchestrating a variety of immune systems, to form an evolutionarily reciprocal and functionally synergistic defense ‘alliance’ against phages.

## Methods

### Bacterial strains and growth conditions

The strains used in this study are listed in Supplementary Data 2*. E. coli* DH5α strain was used for plasmid construction, and *E. coli* BL21 (DE3) strain was used for protein expression and purification. *E. coli* MG1655 was used as the host *E. coli* strain for each phage. All bacterial strains were cultured at 37°C in Lysogeny Broth (LB) media containing 10 g/L tryptone, 5 g/L yeast extract, 10 g/L NaCl. Solid plates were prepared using LB medium supplemented with 15 g/L agar, while liquid cultures were grown with agitation at 200 rpm. Phages T1, T2, T3, T4, T5, T6, T7, and EP02SG were propagated in liquid LB medium. Cultures were centrifuged and the supernatant was filtered through a 0.22-mm filter, then phage titer was determined by the small drop plaque assay. Main chemicals used in this study are listed in Supplementary Data 2.

### Plasmid construction and transformation

The plasmids, oligonucleotides, and synthetic genes used in this study are listed in Supplementary Data 2. The double-stranded DNA fragments were amplified using Phanta Super-Fidelity DNA polymerase (Vazyme Biotech, Nanjing, China), followed by assembly into a predigested plasmid using the Trelief® Seamless Cloning Kit (Tsingke, Beijing, China) by the Gibson assembly strategy. The constructed plasmids were verified by DNA sequencing.

The plasmids were transformed into *E. coli* DH5α, and *E. coli* BL21 (DE3) using commercially available competent cells. Competent *E. coli* MG1655 cells were prepared using the Ultra-Competent Cell Preps Kit (Sangon Biotech. Shanghai, China). Then 200 ng of plasmids were transformed into 100 μl of competent cells. After 1 h recovery culture at 37°C in 500 μl of LB, the culture was serially diluted tenfold with LB and plated on selected plates containing the required antibiotics. Transformation efficiency was calculated by counting the colonies on the plates and multiplying the value by the dilution ratio. Each assay was conducted with three biological replicates.

### RNA extraction and small RNA sequencing

*E. coli* MG1655 log phase cells were harvested for RNA extraction. Total RNA was extracted using the Bacterial RNA Kit (Omega BIO-TEK, USA) according to the manufacturer’s protocol. For small RNA sequencing, 50 μg RNA was treated with T4 polynucleotide kinase (New England Biolabs, MA, USA) at 37°C. The RNA was then purified by the phenol-chloroform method and then precipitated with an equal volume of isopropanol. The resulting RNA sample was re-dissolved in RNA-free water prior to sequencing. A small RNA library was constructed using the NEXTFLEX Small RNASeq Kit (Bioo Scientific, TX, USA), targeting RNA molecules between 30 and 300 nt. The library was then subjected to Illumina HiSeq sequencing (paired-end, 150 bp reads). The resulting raw data reads were adapter removed and were mapped to the *crlRNA* sequence using previously reported Perl scripts.

### Primer extension analysis

Primer extension analysis for *E. coli* MG1655 was performed based on the previously described method^8^, using the 5’-FAM-labelled *gfp*-specific primer as the reverse transcription primer. Briefly, 10 μg of the total RNA from *E. coli* MG1655 was first digested with RQ1 DNase (Promega, WI, USA), followed by reverse transcription to complementary DNA (cDNA) using 30 enzyme units(U) of the Moloney Murine Leukemia Virus reverse transcriptase (MMLV-RT) (Promega, WI, USA). The extension products were then analyzed using the ABI3730xl DNA Analyzer (Thermo Fisher Scientific, MA, USA), and the results were viewed using Peak Scanner Software v1.0.

### Efficiency of plaque assays

For the experiments shown in Fig. 1d, Fig. 2c, Fig. 3, Fig. 5c, and Extended Data Fig. 6, phage stocks were serially diluted 10-fold in LB and the dilutions were plated onto double layer agar plates of *E. coli* MG1655 with empty vector or defense systems. In Fig. 4a, 2 μL of each dilution was spotted onto double layer agar plates of *E. coli* MG1655.

### Infection dynamics in liquid culture

180 μL of early-log cultures were transferred into wells in 96-well plates containing 20 μL of phage diluent for a final MOI of 0.01/0.04 or 10 for phage T4 and λ, or 20 μL of LB medium for uninfected control. Infections were performed in triplicate from cultures prepared from three separate colonies. OD_600_ values were measured every fifteen minutes for six hours in a real-time plate reader Stratus 600 nm (Cerillo, Charlottesville, VA, USA) by continuously shaking the culture plate at 37°C.

### Fluorescence measurement

*E. coli* MG1655 cells expressing *gfp* were grown until the late exponential phase, and the OD_600_ and fluorescence were measured simultaneously using the Synergy H4 Hybrid multimode microplate reader (BioTeck, VT, USA). At least three individual colonies were randomly selected and cultured for each experimental setting. The fluorescence/OD_600_ ratio was calculated for each of the three individual biological samples, and the mean and standard deviation were determined accordingly.

### Protein expression and purification

Then plasmids pET28a-T7-cas5c and pET28a-T7-Erlang were transformed into *E. coli* BL21 (DE3) to express Cas5C and ErlA-ErlB complex, respectively. *E. coli* BL21 was then inoculated into 1 L LB medium (50 μg/mL kanamycin) and cultured on a shaker at 37°C. Cultures were grown to an optical density (OD_600_) of 0.6-0.8 and then induced with 0.1 mM IPTG for 8 h at 22°C. Protein purification was performed on a His-Trap HP column (Cytiva) at 4°C according to standard protocols. Harvested cells were collected by centrifugation (2350 × g, 30 min, 4°C) and resuspended in 30 mL of binding buffer (20 mM HEPES pH = 7.5, 0.5 M NaCl, 20 mM imidazole, 5% glycerol). The cells were then sonicated on ice and after centrifuged (15000 g, 1 h, 4°C) to obtain the supernatant. The supernatant was applied to His-Trap HP column equilibrated with the binding buffer and washed with the same buffer containing 20 mM imidazole. Proteins were eluted with the binding buffer containing 250 mM imidazole. Affinity-purified proteins were fractionated and identified by SEC using a GE Life Sciences Superdex 200 10/300 column. The protein concentration was determined by the BCA method using bovine serum albumin (BSA) as a standard. The purified protein was then stored at -80°C.

### RNA cleavage assays

RNA used in the cleavage assays is listed in Supplementary Data 2. The cleavage assay was performed in 1× cleavage buffer [20 mM HEPES-NaOH (pH 7.5), 100 mM KCl, 5% glycerol, and 1 mM DTT]. 2.5 μM Cas5c was incubated with 0.25 μM 5′ FAM labeled RNA at 37°C for 30 min. The reaction products were then mixed with formamide-containing dye and analyzed on 15% urea PAGE. The urea PAGE gels were analyzed on a Tanon-5200 Multi instrument (Tanon Science & Technology Co. Ltd., Shanghai, China).

### Stability of Acr-expressing plasmids in the *E. coli* MG1655 cell population

300 ng plasmids expressing *gfp* and 300 ng plasmids expressing *acr* and *mcherry* were mixed and transformed into *E. coli* MG1655. After incubation in LB medium for 1 h, the culture was inoculated into fresh LB medium (containing Str and Kan) with 1%, and fluorescence was measured after 10 h of incubation. This procedure was repeated four times, with the addition of 0.1 mM IPTG on the second, third, and fourth occasions.

### DNA cleavage assay

Phage genomic DNA was extracted using the Lambda phage Genomic DNA Kit (Zoman Biotechnology Co., Ltd., Beijing, China) according to the manufacturer’s protocol. For DNA cleavage assay, a final concentration of 1 mM protein and 200 ng DNA substrate were incubated in reaction buffer (20 mM HEPES pH 8.0, 25 mM NaCl, and 5 mM MgCl_2_) at 37°C for 2 h. The reactions were stopped by adding 10 mM EDTA and deproteinized with Proteinase K (100 μg/mL) for 20 min at 37°C. Samples were separated in 1% agarose gel electrophoresis, and the results were visualized using the Chemical Imaging System (Bio-Rad).

### NADase activity assay

50 μM Sole-Sir2 and 2 mM NAD were incubated in 50 μL Tris-HCl buffer (50 mM, pH 7.5). Reactions were performed at 25 °C and quenched with methanol. The protein precipitate was removed by centrifugation and then the supernatant was analyzed by LC-MS. LC-MS analysis was performed on an AGILENT-1200 HPLC/6520 QTOFMS (USA) system using a C18 analytical column (Ultimate® AQ-C18 250 × 4.6 mm, particle size 5 μm). The mobile phase consisted of water containing 10 mM ammonium acetate (solvent A) and acetonitrile (solvent B). The gradient was as follow: 0-7 min, 98% A; 12 min, 40% A; 15 min, 40% A; 18 min, 98% A; 21 min, 98% A. Flow rate: 1 mL/min.

### Bioinformatic search of CRISIS genes

To explore CRISIS genes, we collected all the annotated Cas5c proteins achieved in the NCBI database, and retrieved their corresponding genome sequences. By manually examining their intergenic sequences, we further screened for SRUs and their target sites. SRUs were predicted based on a palindromic sequence (forming a stem-loop RNA structure) within ΨR and the conserved 7 nt at its 3’ terminal, while target sequences were predicted based on its complementarity to the ΨS sequence of SRU and a flanking PAM motif.

### Preliminary estimation of the frequency of immune systems

To preliminarily estimate the occurring frequency of each defense system in microbial genomes, we downloaded genomic data of bacteria from NCBI in 2020 and constructed a representative dataset by selecting one genome per species (n = 45248 genomes). For each system, initial protein sequences taken from experimentally validated loci were aligned using MAFFT (version v7.487) software and converted into HMM profiles. We used HMMER (version 3.4) to search each protein in the representative dataset against all HMM profiles, applying strict filtering parameters (expect threshold of 1e-8) to minimize the inclusion of unrelated proteins. The resulting alignments were utilized to create a new set of signature gene profiles to the second iteration. For multi-protein systems, a putative locus was considered as a hit if every signature gene profile for the system had a match within the locus, with the maximum distance between signature genes should be less than 3 gene intervals. For single-gene systems, a locus was considered as a hit if the protein matched the single signature gene profile of the system with an expect threshold of 1e-8.

### Bioinformatic analysis

RNA secondary structures were predicted using the RNAfold webserver. Promoter elements were predicted using the BPROM program (Softberry tool). Structures were predicted using AlphaFold3 Server (https://golgi.sandbox.google.com/). Domain analysis was performed using HHpred (https://toolkit.tuebingen.mpg.de/tools/hhpred).

### Statistics & Reproducibility

All experiments in the present manuscript were repeated at least three times independently with similar results.

### Reporting Summary

Further information on research design is available in the Nature Portfolio Reporting Summary linked to this article.

### Data Availability

All relevant data are included in the paper and/or its supplementary information files. Source data are provided as a Source Data file. The raw data for the RNA-seq experiments in Fig. 1, Extended Data Fig. 2, and Extended Data Fig. 3 have been deposited to the National Center for Biotechnology Information (NCBI) with the BioProject accession number PRJNA1159167. All strains and plasmids are available from the corresponding author upon request; requests will be answered within 2 weeks. Source data are provided with this paper.

## Acknowledgements

We thank Prof. Jie Feng for providing the EP02SG phage as a gift. This work was supported by the Strategic Priority Research Program of the Chinese Academy of Sciences [XDB0830000], the National Natural Science Foundation of China [32150020, 32270092, 32230061, 32370090, 32200057, and 32022003], the Youth Innovation Promotion Association of CAS [2020090], and the China National Postdoctoral Program for Innovative Talents [BX20220331].

## Author Contributions

M.L. conceptualized this study. M.L., X.S., R.W., designed the experiments. X.S. and R.W. constructed the plasmids. F.C., R.W., A.W., and X.S. conducted the efficiency of plaquing assays. X.S. and R.W. performed the fluorescence measurement and bacteria transformation assays with the assistance of Q.X. X.C. and C.L.. F.C. and X.S. performed RNA-seq and primer extension. X.S. and M.L. performed the bioinformatic analyses. Z.L. conducted protein purification and liquid culture growth assays. Formal analysis of results was done by M.L., X.S., R.W., and Y. F.. M.L. wrote the paper, which was edited and approved by all authors.

## Competing Interests

M. L., X. S., and R. W. filed a related patent.

## Additional information

**Supplementary Information** is available in the online version of the paper.

**Correspondence and requests for materials** should be addressed to M. L.

**Reprints and permissions information** is available at www.nature.com/reprints.

## Extended Data

**Extended Data Figure 1:**
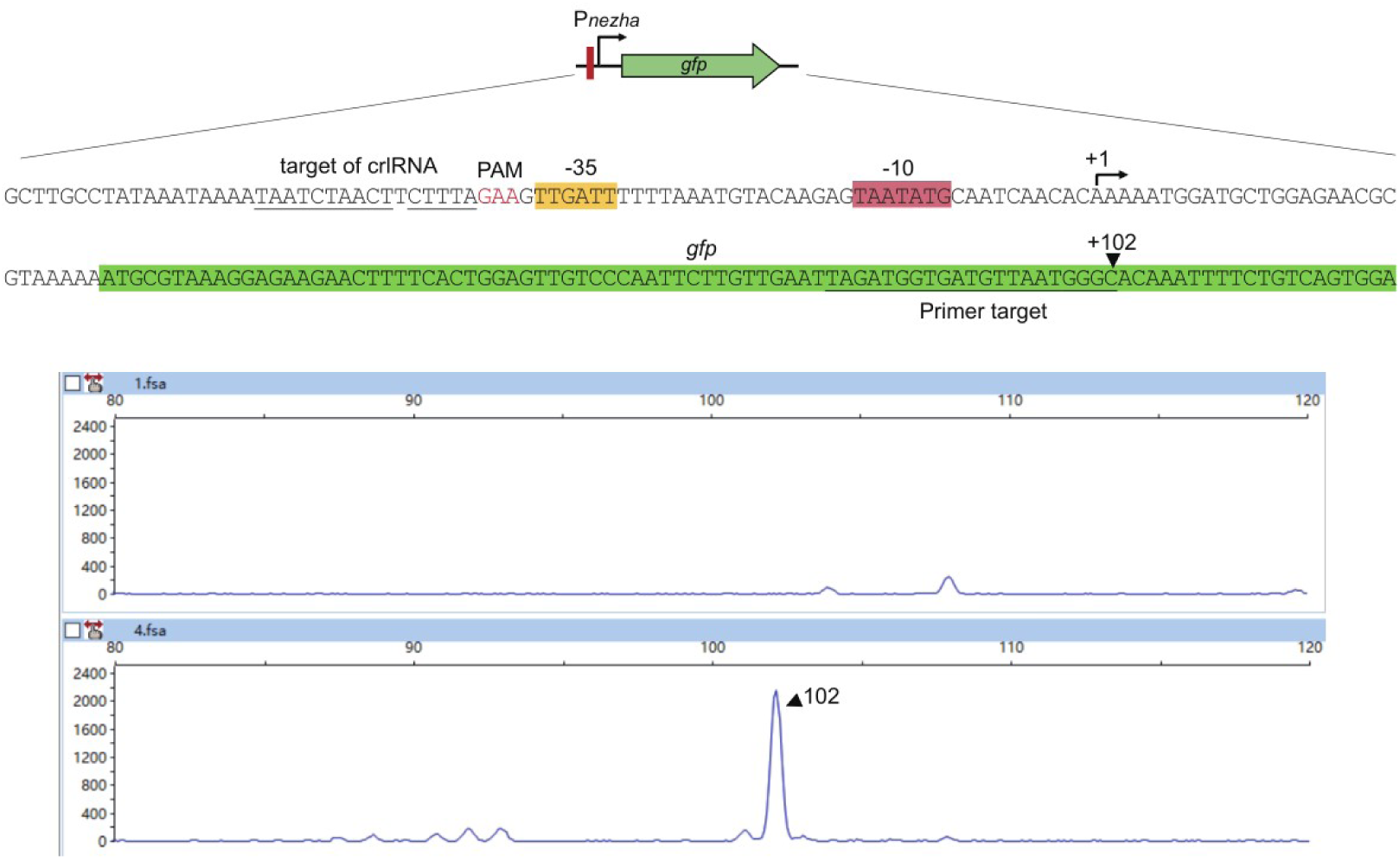
Primer extension assay to determine the TSS of the Nezha system from *N. subflava* CCUG 29761. P*_nezha_* was first linked to a *gfp* gene and then introduced into *E. coli* MG1655 cells. Total RNA was extracted from the cells for primer extension analysis. The primer used for the assay was designed to target the *gfp* RNA transcript and labelled with 5′-FAM (see Methods section for details). The resulting cDNA products from the primer extension assay were analyzed for fragment sizes.

**Extended Data Figure 2:**
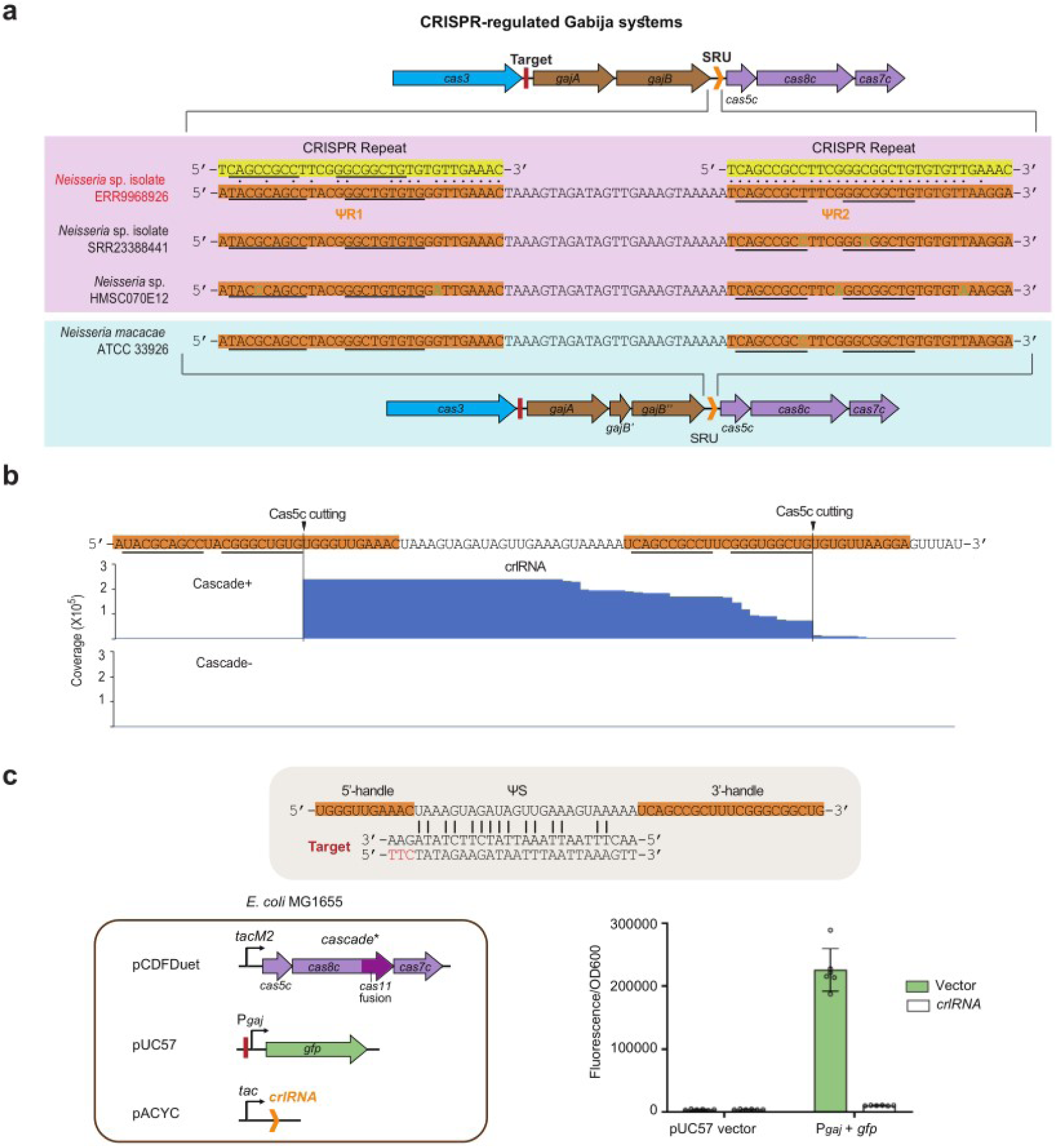
Characterization of a CRISPR-regulated Gabija system. **a**, Genetic organization of CRISPR-regulated Gabija systems from four *Neisseria* species. Dots indicate identical nucleotides shared between the CRISPR repeat and the repeat-like sequences (ΨR1 and ΨR2) of SRUs (solitary repeat units). Palindromic nucleotides are underlined. The SRU from *Neisseria* sp. isolate ERR9968926 (highlighted in red) was experimentally characterized in panels **b** and **c**. For other SRUs, nucleotides differing from the experimentally characterized SRU are indicated in green. **b**, Determination of the regulatory crlRNA produced from the SRU by sRNA-seq. Processing sites of Cas5c are indicated. Total RNA samples from *E. coli* cells encoding (Cascade+) or lacking (Cascade-) the Cascade proteins were separately subjected to sequencing. **c**, Fluorescence from a *gfp* gene controlled by the promoter of Gabija system (P*_gaj_*) in the presence of the crlRNA and Cascade proteins. The complementarity between the P*_gaj_* and crlRNA is depicted, with PAM nucleotides highlighted in red. The red bar indicates the target site of crlRNA. *tacM2*, a mutated version of *tac* promoter. The *cas11* gene is fused within *cas8c*. Data are presented as the mean ± s.d. from six biologically independent replicates. Note that *N. subflava* CCUG 29761 Cas proteins regulating the Nezha system were used in panels **b** and **c** for convenience.

**Extended Data Figure 3:**
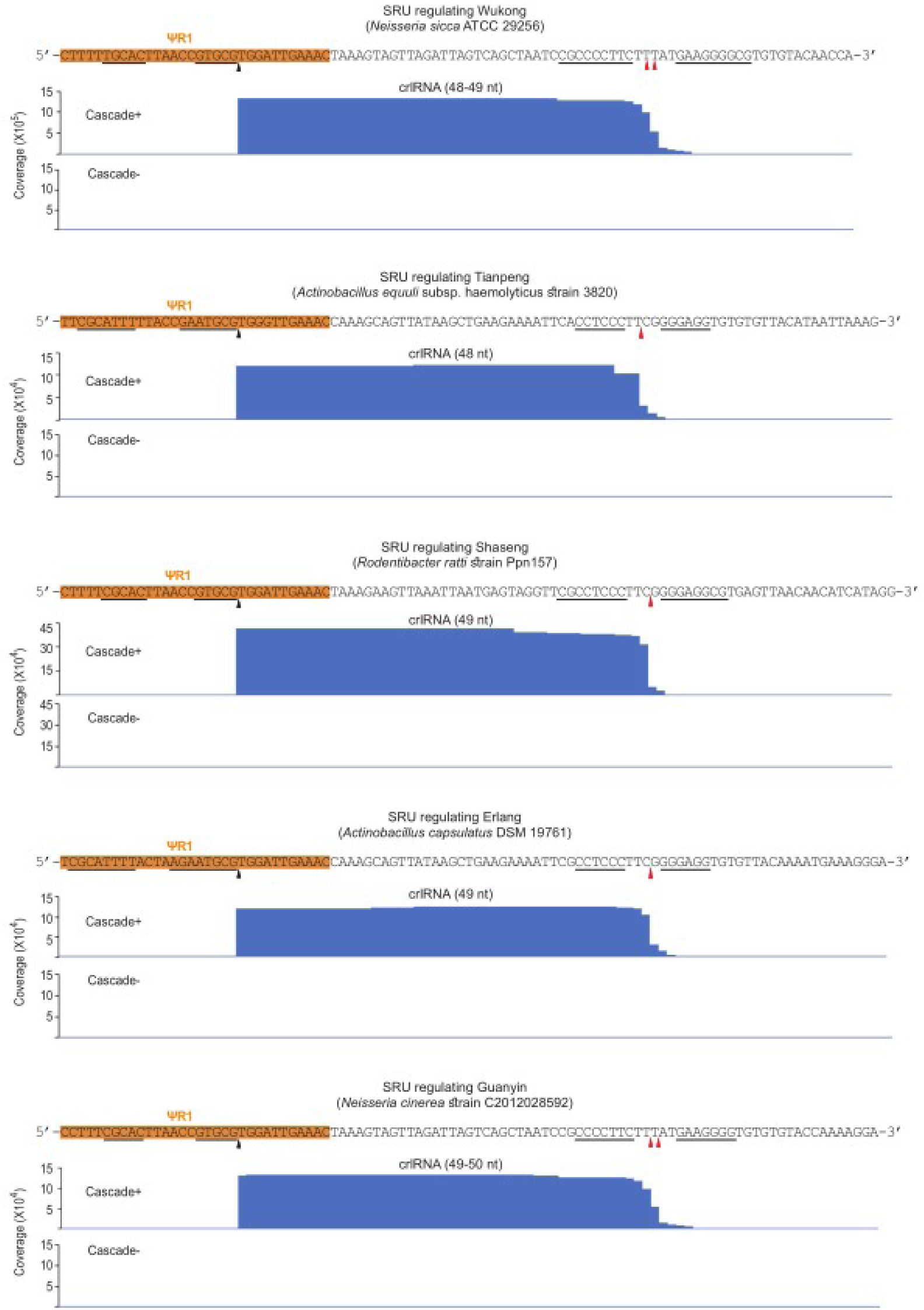
sRNA-seq to determine the sequence of crlRNAs regulating different CRISIS systems. Palindromic nucleotides that form the stem of a hairpin RNA structure are underlined. The black triangles indicate the canonical RNA processing site of Cas5c, while the red triangles indicate flexible 3′ termini within the loop of a 3′ hairpin structure. The sizes of primary crlRNA products are given in brackets. Note that total RNA samples from *E. coli* cells encoding (Cascade+) or lacking (Cascade-) the Cascade proteins regulating the Nezha system were separately subjected to sequencing.

**Extended Data Figure 4:**
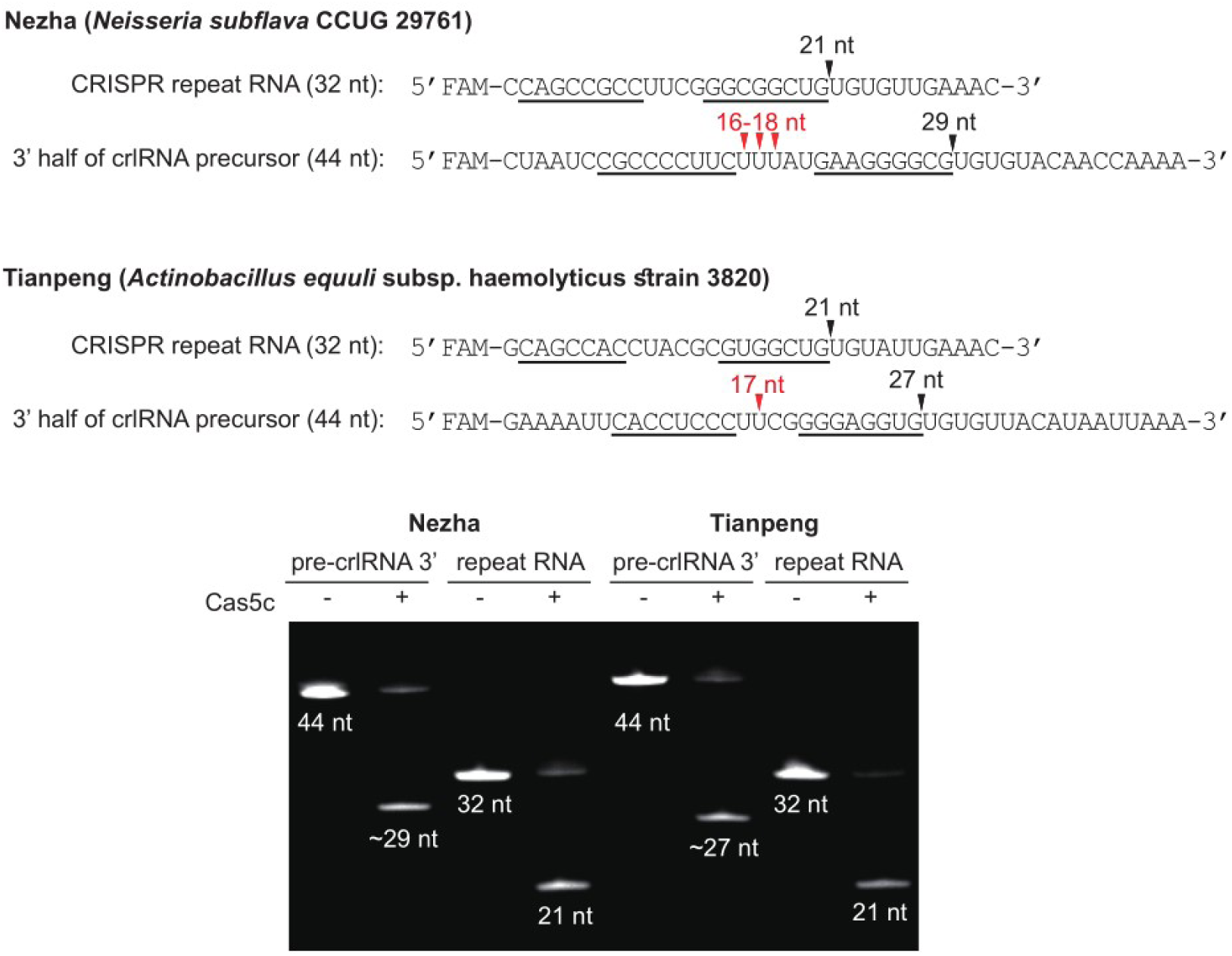
Processing site of Cas5c on the 3′ hairpin structure of crlRNA precursors (pre-crRNA) associating with Nezha and Tianpeng. Oligoribonucleotides are 5′ FAM-labelled and digested by Cas5c *in vitro*. The conserved CRISPR repeat RNA from the CRISPR-Cas systems regulating Nezha or Tianpeng was also included for comparison. Palindromic nucleotides forming the stem of the hairpin structure are underlined. Black triangles indicate the processing sites of Cas5c, while red ones indicate the 3′ termini of the corresponding mature crlRNAs determined by sRNA-seq (Figure 1a and Extended Data Fig. 3).

**Extended Data Figure 5:**
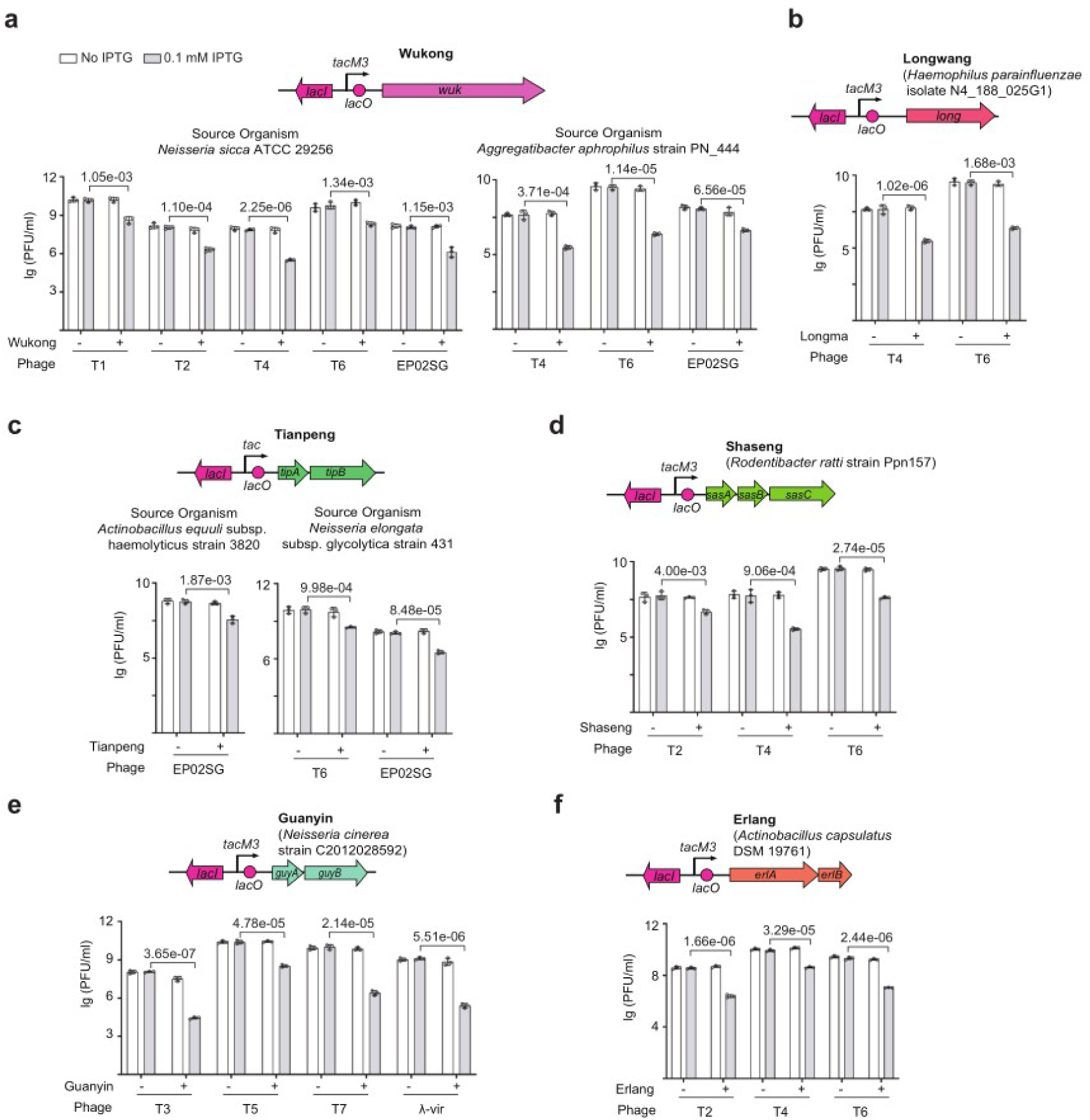
The immunity of various CRISIS systems against a series of *E. coli* phages. Each CRISIS system was placed under the control of a synthetic promoter (*tac* or *tacM3*) containing the *lacO* operator, and introduced into *E. coli* MG1655 cells using the pET-28a vector to assess its phage immunity. Data are presented as the mean ± s.d. from three biologically independent replicates.

**Extended Data Figure 6:**
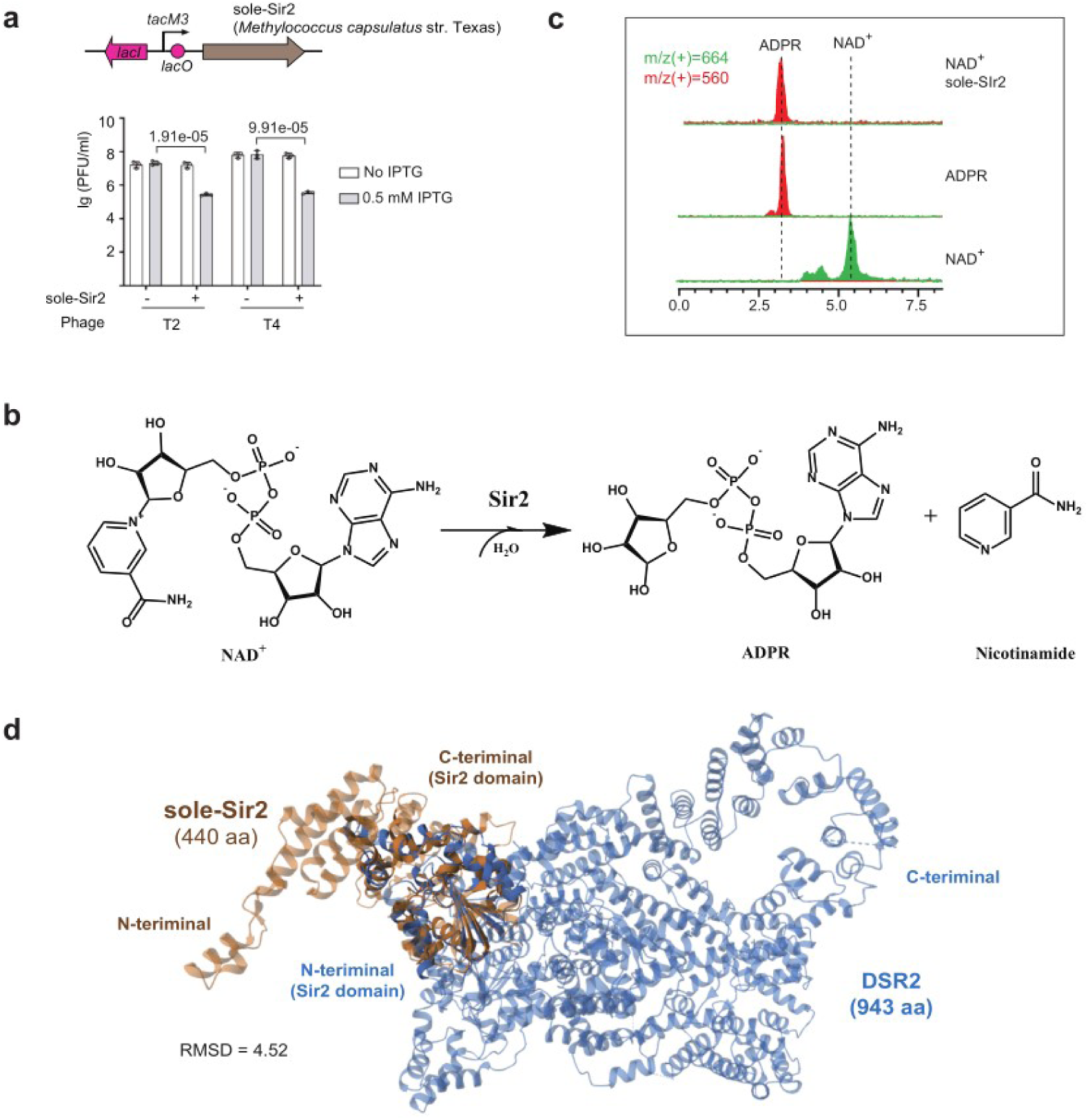
Characterization of a homolog of the CRISPR-regulated sole-Sir2. **a**, Assessment of immunity of sole-Sir2 against T2 and T4 phages. Note that this sole-Sir2 protein is not regulated by CRISPR-Cas, but is a close homolog of those listed in Fig. 2c. This protein was placed under the control of an IPTG-inducible *tacM3* promoter and expressed from pET-28a. Data are presented as the mean ± s.d. from three biologically independent replicates. **b**, Illustration of the canonical chemical reaction catalyzed by Sir2-containing proteins. **c**, LC-MS assay to evaluate the catalytic activity of sole-Sir2. **d**, Structural prediction and alignment with the recently characterized DSR2 protein from *Bacillus subtilis* 29R (Zhang JT, et al., 2024). The RMSD value was calculated based on a separate alignment of the Sir2 domains of sole-Sir2 and DSR2.

**Extended Data Figure 7:**
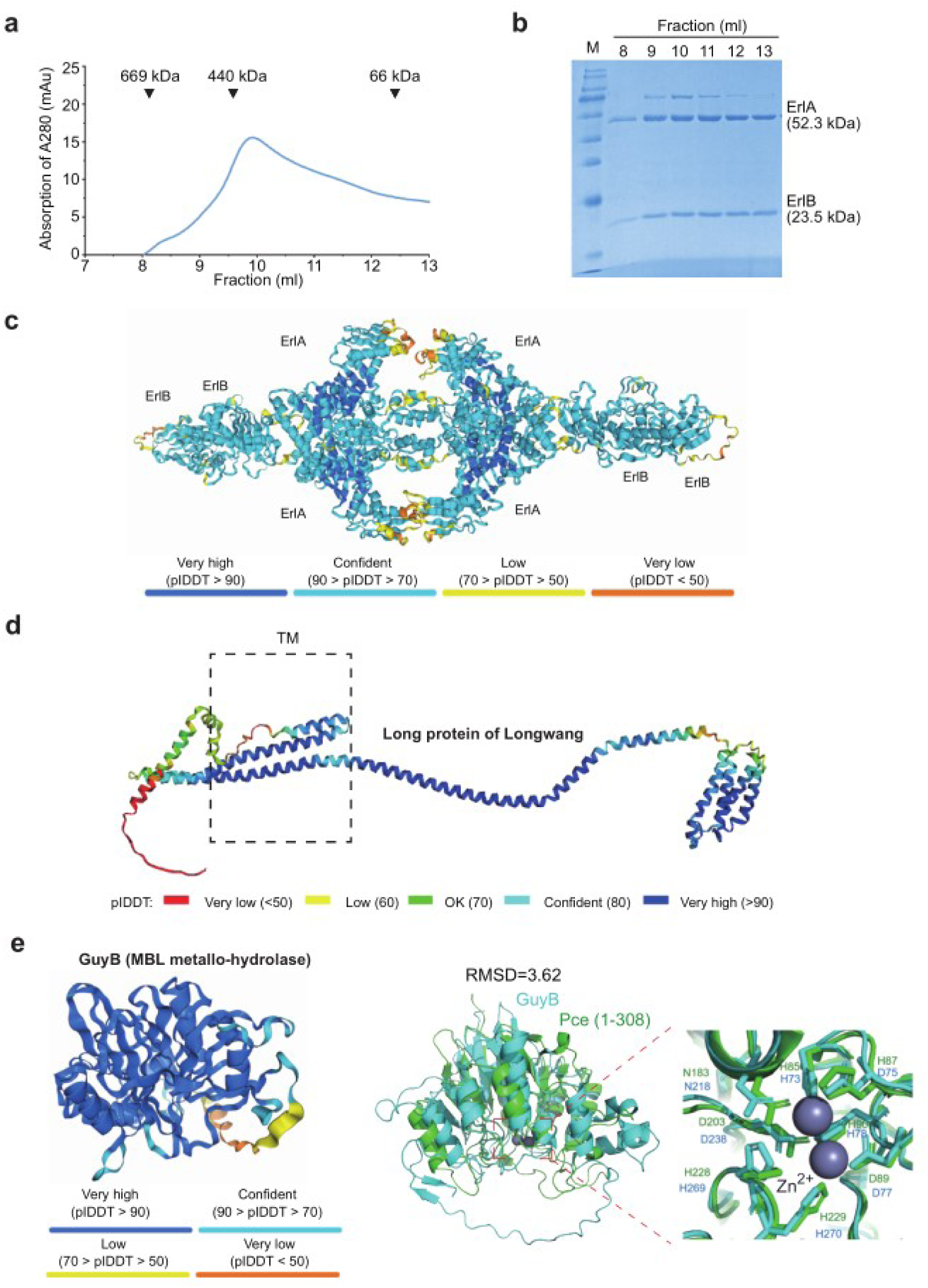
Biochemical and structural analysis of CRISIS effector proteins. **a-c**, Purification of the Erlang complex (composed of ErlA and ErlB) and prediction of its structure. Standard proteins are labelled on the chromatogram to estimate the molecular weight of the Erlang complex. kDa, kilodaltons. **d**, Structural prediction of the Long protein from the Longwang system. TM, transmembrane helices. **e**, Structural prediction of GuyB (the MBL metallo-hydrolase encoded by Guanyin system) and its resemblance to the N-terminal Zn^2+^ catalytic module of Pce protein (Juan A Hermoso et al., 2005).

**Extended Data Figure 8:**
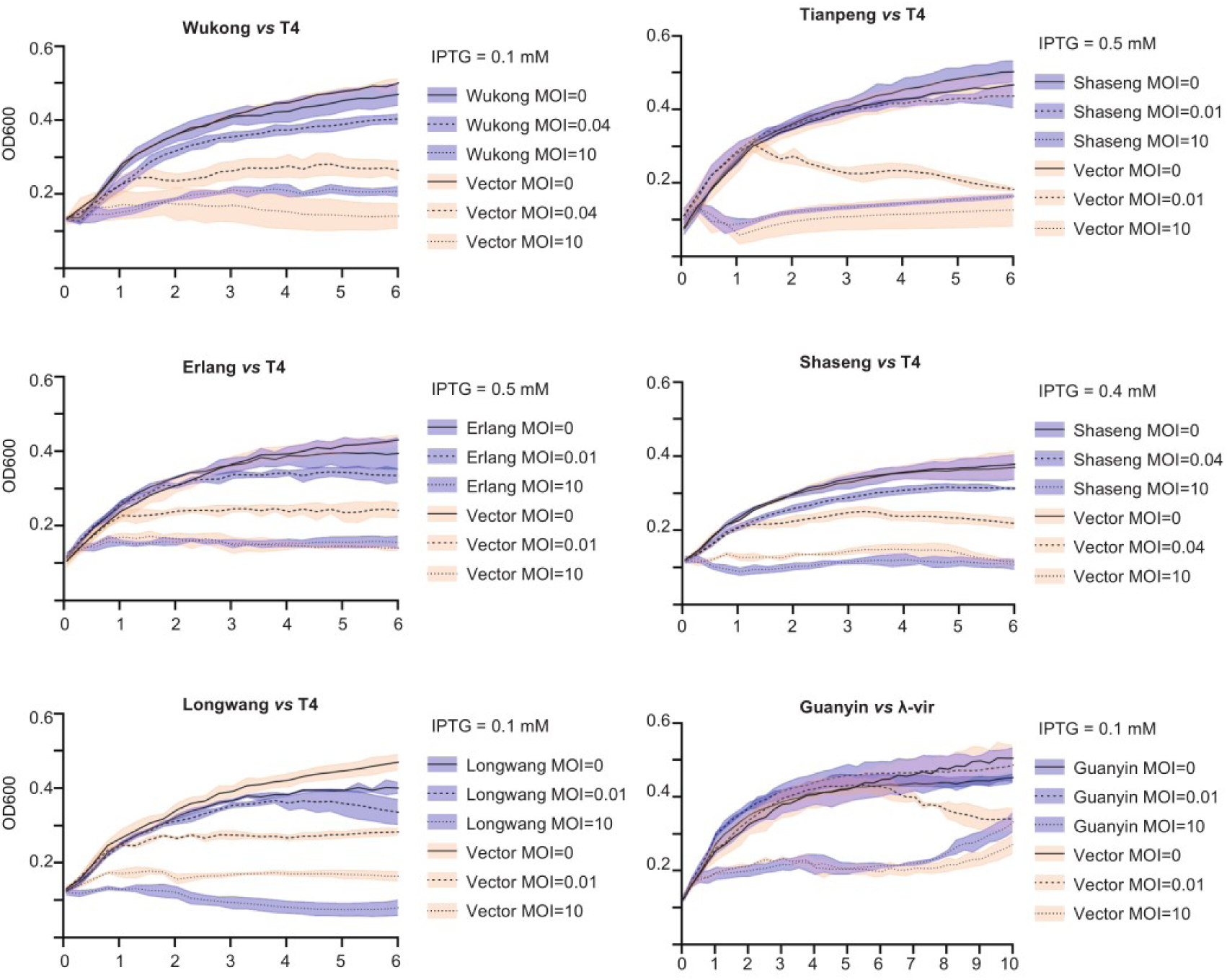
Abi responses induced by CRISIS systems. Growth curve of *E. coli* cells encoding different CRISIS systems with or without the infection by T4 or the virulent λ phage under different MOIs. Data are presented as the mean ± s.d. from three biologically independent replicates.

**Extended Data Figure 9:**
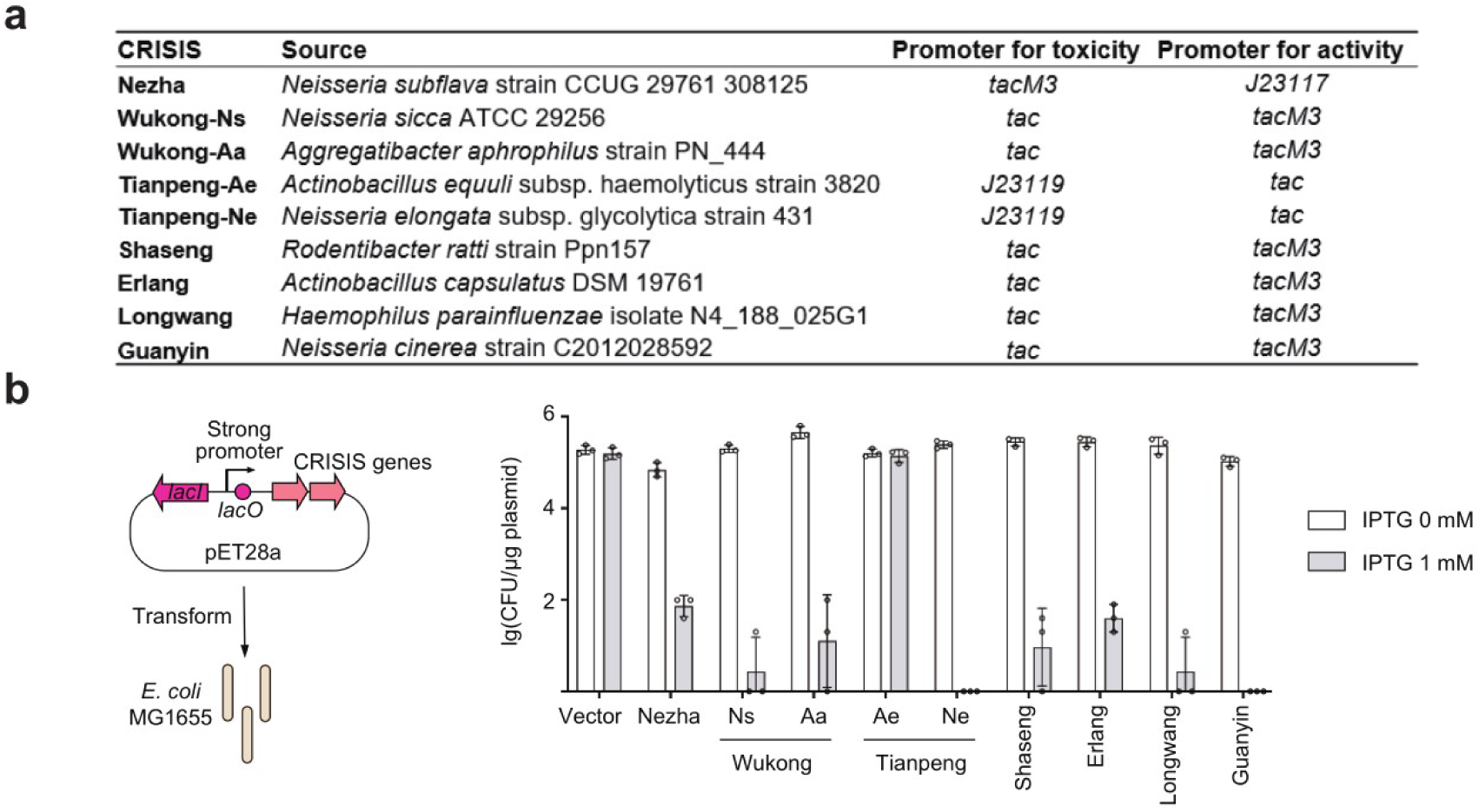
Cytotoxicity upon the overexpression of CRISIS systems. **a**, The different promoters used to evaluate the toxicity and the immune activity of CRISIS systems, respectively. The source organism of each tested CRISIS system is given. **b,** Transformation efficiency of *E. coli* cells by a pET28a derivate over-expressing different CRISIS genes. Data are presented as the mean ± s.d. from three biologically independent replicates.

**Extended Data Figure 10:**
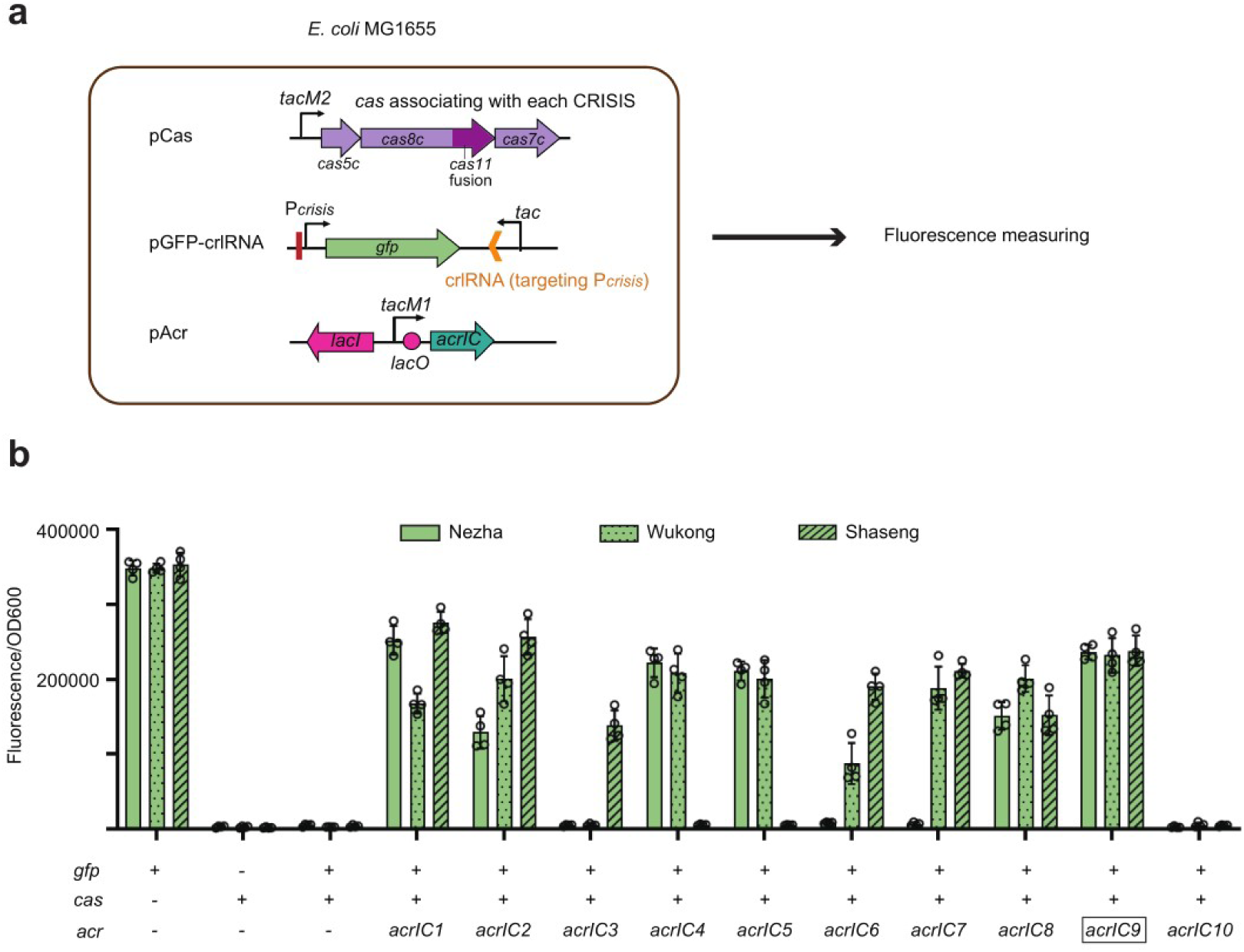
Screening for Acr proteins that inhibit the Cascade proteins regulating Nezha, Wukong, or Shaseng. **a,** Scheme illustrating the experimental design. **b,** Fluorescence from the *gfp* gene under the promoter of each CRISIS system in *E. coli* cells expressing their corresponding crlRNAs and Cascade proteins, as well as one of AcrIC1-AcrIC10. The AcrIC9 protein was selected for the assays in Fig. 4 and 5. Data are presented as the mean ± s.d. from four biologically independent replicates.

**Extended Data Figure 11:**
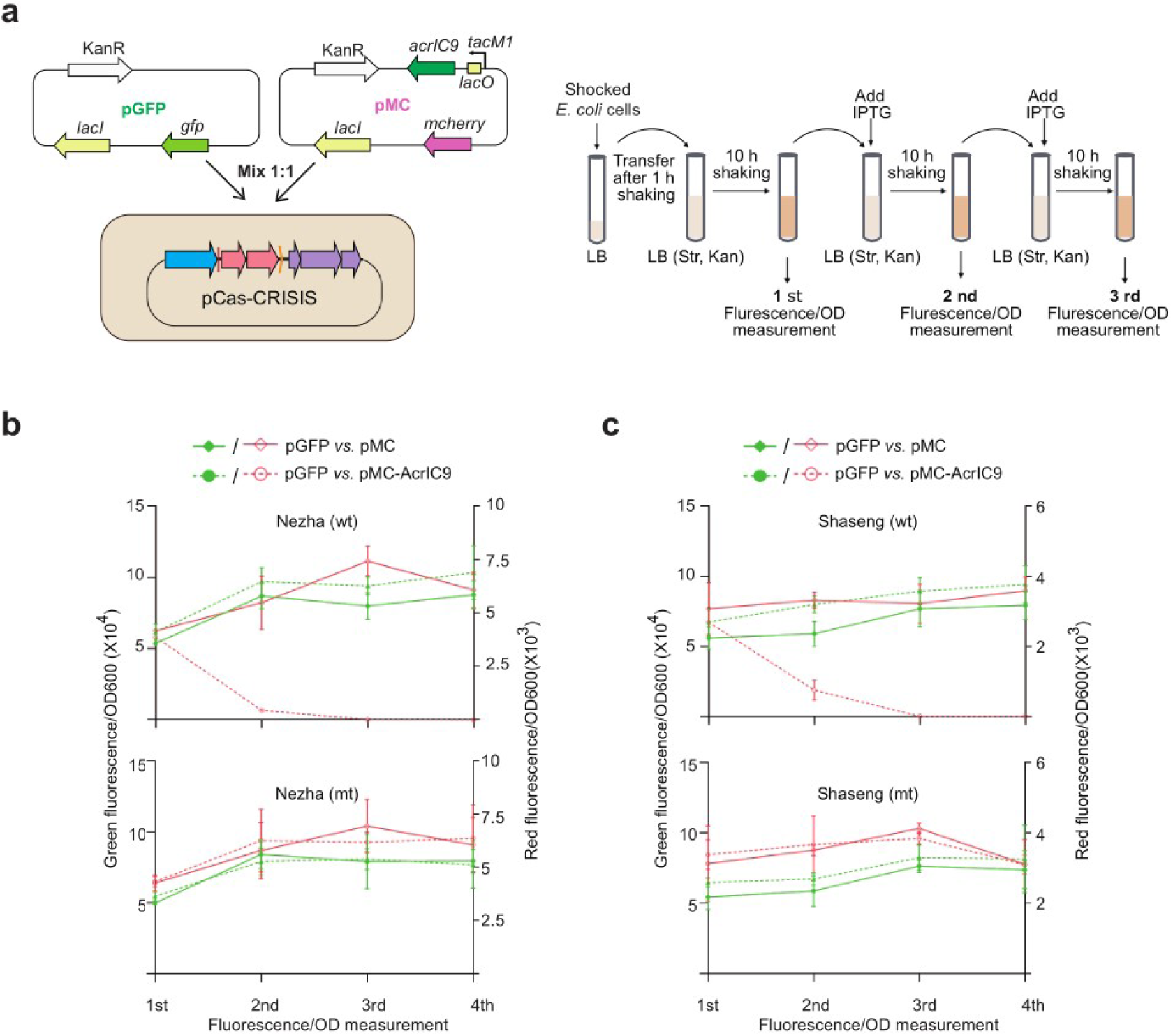
The anti-anti-CRISPR effect of Nezha (b) and Shaseng (c). **a,** Illustration of the experimental procedure. **b-c**, The persistence of an AcrIC9-encoding plasmid (pMC-AcrIC9; also carrying the *mcherry* reporter gene) in the population of *E. coli* cells containing pCas-Nezha or pCas-Shaseng (or their derivatives with mutated CRISIS genes). A competing plasmid (with the same backbone) carrying *gfp* (pGFP) was mixed with pMC-AcrIC9 or pMC (lacking the *acrIC9* gene) at a 1:1 ratio before co-transformation into *E. coli* cells. Data are presented as the mean ± s.d. from three biologically independent replicates.

**Extended Data Figure 12:**
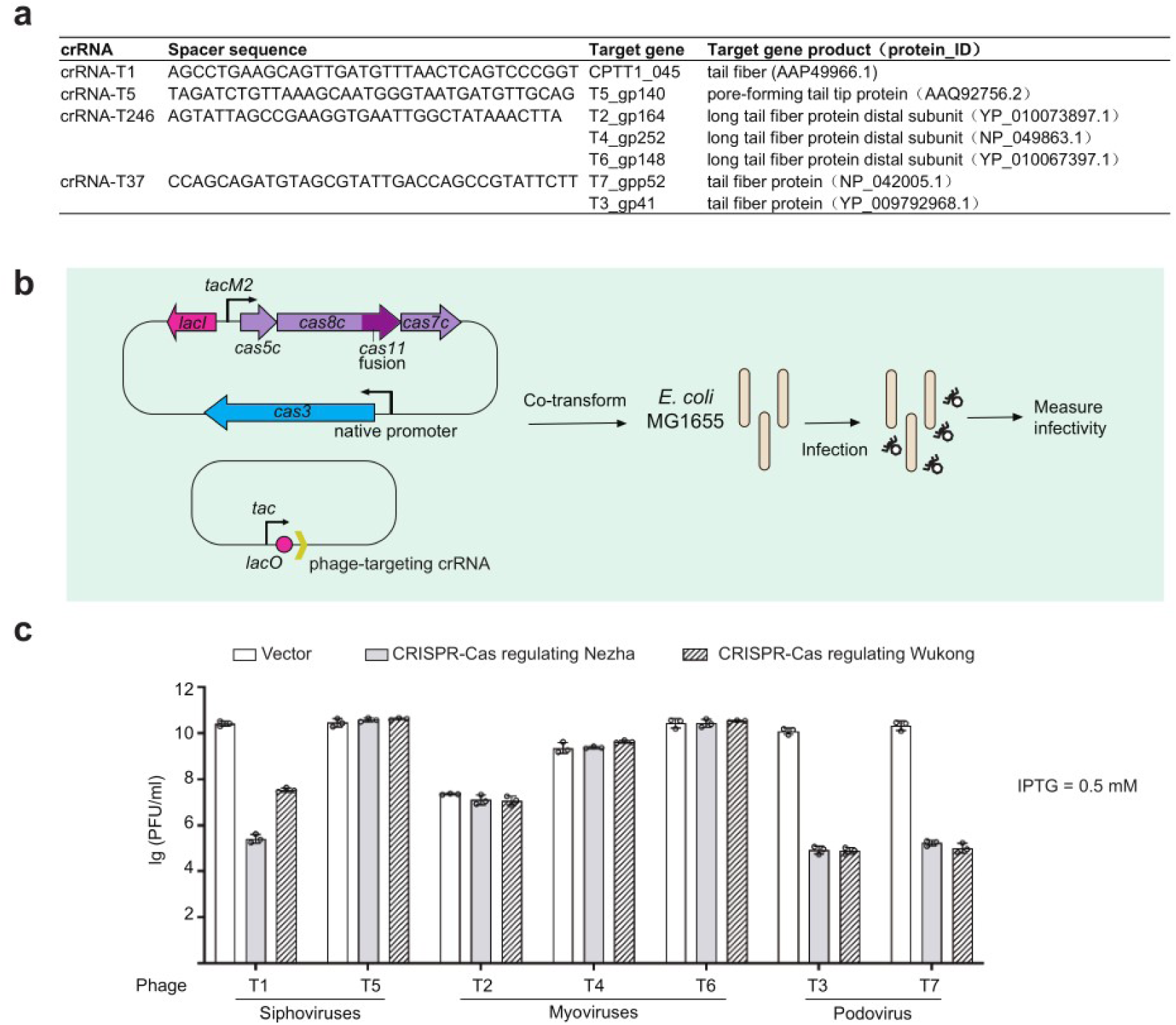
Anti-phage activity of the CRISPR-Cas systems supervising Nezha and Wukong. **a,** Design of phage-targeting crRNAs. The spacers target phage sequences downstream of a 5′-TTC-3′ PAM sequence. **b,** Scheme illustrating the plasmid design and the experimental procedure. The *cas3*, *cas5c*, *cas8c* (containing a fused *cas11*) and *cas7c* genes associating with Nezha or Wukong were cloned into pCDFduet. Phage-targeting crRNAs were expressed under an inducible *tac* promoter from a pACYC derivate. **c,** Phage immunity conferred by the CRISPR-Cas system governing Nezha or Wukong. Data are presented as the mean ± s.d. from three biologically independent replicates.

**Extended Data Figure 13:**
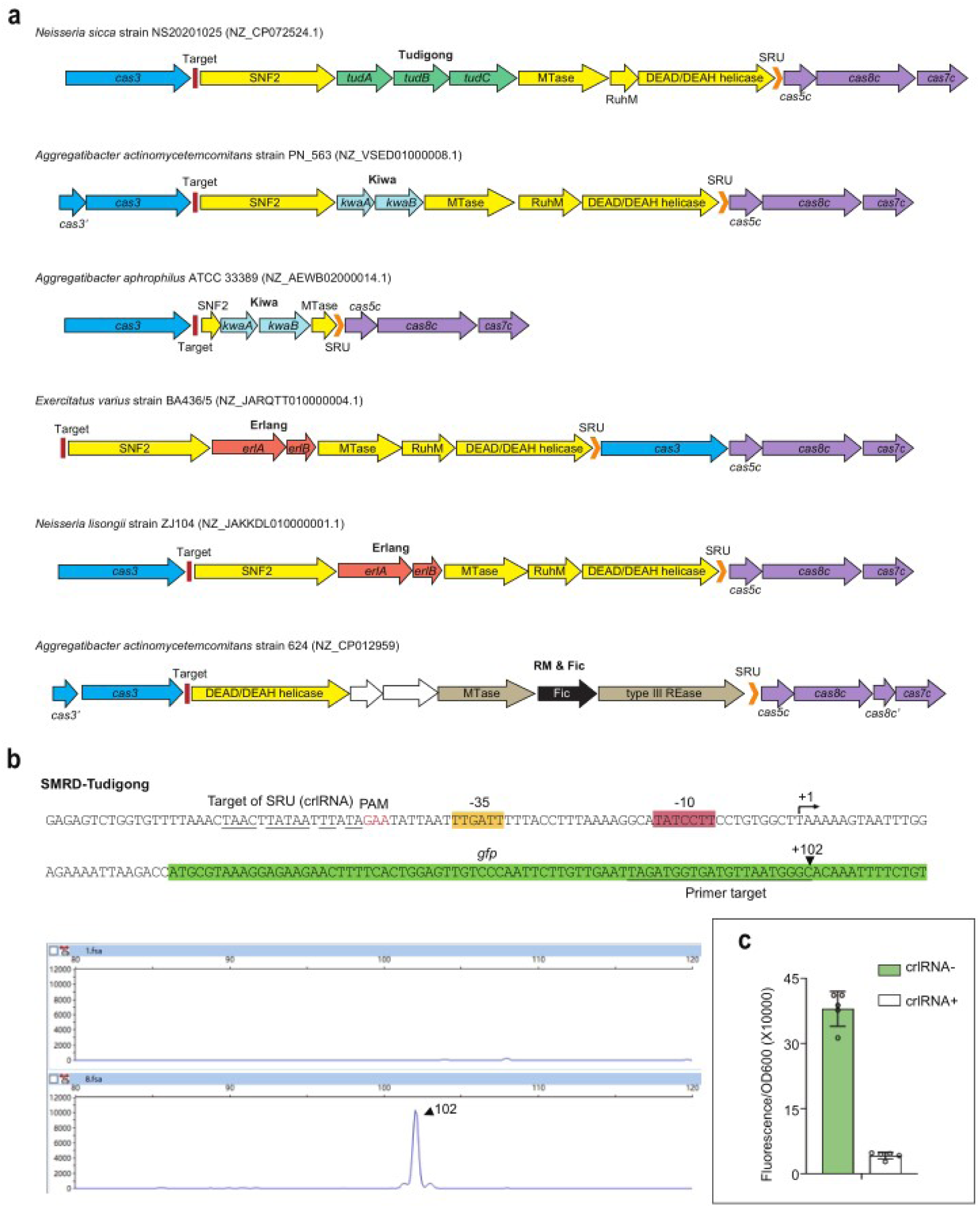
Characterization of CRISPR regulation on the SMRD defensive gene cluster. **a**, Scheme depicting the architecture of CRISPR-regulated SMRD gene clusters, which preferentially encoding a SNF2-related protein, a DNA methyltransferase, a RuhM family nuclease, and a DEAD helicase, as well as a known defense system (such as Tudigong, Kiwa, or Erlang). **b**, Determination of the TSS of the SMRD cluster containing a Tudigong system (designated as SMRD-Tudigong; from *N. sicca* strain NS20201025). Predicted promoter sequence was linked to *gfp* and introduced into *E. coli* MG1655 cells, from which total RNA was extracted for primer extension analysis. The primer used for the assay was designed to target the *gfp* RNA transcript and was labelled with 5′-FAM. The resulting cDNA products from the primer extension assay were analyzed for fragment sizes. **c,** Fluorescence from *gfp* controlled by the promoter of SMRD-Tudigong in *E. coli* cells expressing or lacking the corresponding crlRNA. Note that the *N. subflava* CCUG 29761 Cas proteins regulating Nezha were also expressed in these cells. Data are presented as the mean ± s.d. from five biologically independent replicates.

**Extended Data Figure 14:**
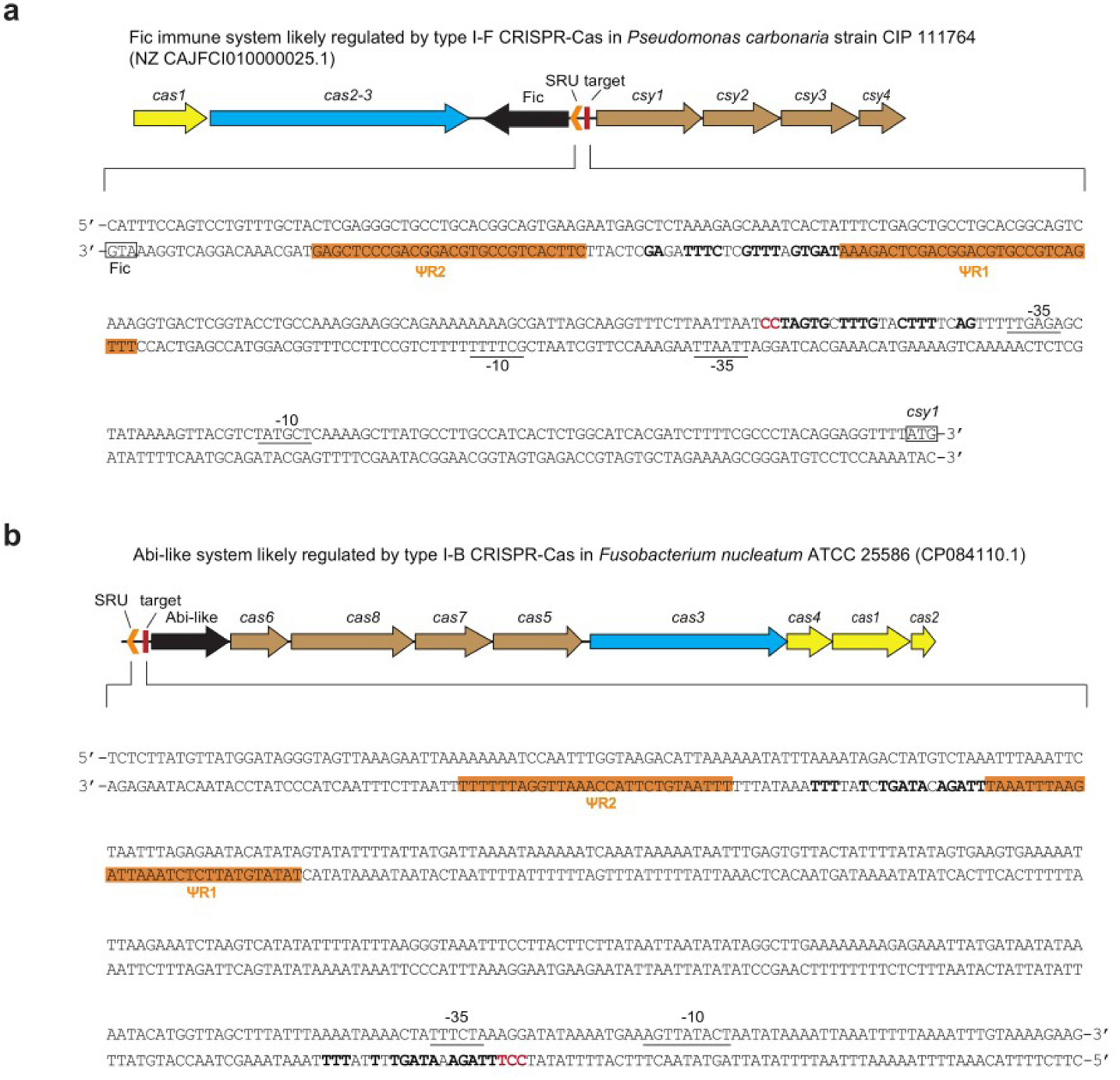
Immune systems potentially regulated by I-F (a) and I-B (b) CRISPR-Cas systems. The repeat-like sequences (ΨR1 and ΨR2) of each SRU are indicated. The identical nucleotides shared between the spacer-like (ΨS) sequence of each SRU and its target protospacer are highlighted in bold. PAM nucleotides are shown in red. Predicted promoter elements (-10 and -35) are underlined. Start codons are framed.

